# Gemcitabine and ATR inhibitors synergize to kill PDAC cells by blocking DNA damage response

**DOI:** 10.1101/2024.03.22.586243

**Authors:** Stefanie Höfer, Larissa Frasch, Kerstin Putzker, Joe Lewis, Amirhossein Sakhteman, Matthew The, Florian P. Bayer, Julian Müller, Firas Hamood, Bernhard Kuster

**Affiliations:** Chair of Proteomics and Bioanalytics, Technical University of Munich, Freising, Germany; Chemical Biology Core Facility, EMBL Heidelberg, Heidelberg, Germany; German Cancer Consortium (DKTK), Partner Site Munich, Munich, Germany

## Abstract

The DNA-damaging agent gemcitabine (GEM) is a first-line treatment for pancreatic cancer but chemoresistance is frequently observed. Several clinical trials investigate the efficacy of GEM in combination with targeted drugs including kinase inhibitors but the experimental evidence for such rational is often unclear. Here, we phenotypically screened 13 human pancreatic adenocarcinoma (PDAC) cell lines against GEM in combination with 140 clinical kinase inhibitors and observed strong synergy for the ATR inhibitor Elimusertib in most cell lines. Dose-dependent phosphoproteome profiling of four ATR inhibitors following DNA damage induction by GEM revealed a strong block of the DNA damage response pathway including phosphorylated pS468 of CHEK1 as the underlying mechanism of drug synergy. The current work provides a strong rationale for why the combination of GEM and ATR inhibition may be useful for the treatment of PDAC patients and constitutes a rich phenotypic and molecular resource for further investigating effective drug combinations.

## Introduction

Pancreatic ductal adenocarcinoma (PDAC) is a devastating disease and, unlike for other malignancies, survival rates have barely increased over the past decades^1–3^. Patients diagnosed with early, localized PDAC typically undergo surgical resection, often complemented by chemotherapy or radiotherapy to enhance disease control. Unfortunately, the majority of pancreatic tumors are diagnosed late, rendering surgical intervention ineffective and, instead, chemotherapy is the primary systemic treatment modality^4^. One frontline drug is Gemcitabine (GEM), a nucleoside analogue that induces DNA damage and replication stress response in dividing cells by disrupting DNA synthesis^4–6^. In PDAC, GEM is often administered in combination with albumin-bound paclitaxel particles (nab-paclitaxel) as the combination substantially improves therapeutic efficacy^7^. Another standard therapy is FOLFIRINOX, a multi-chemodrug regimen including folinic acid, 5-fluorouracil, irinotecan, and oxaliplatin^8^. While FOLFIRINOX can achieve superior therapeutic outcomes, it also comes with higher toxicity, which limits its use to patients with good overall performance status^8,9^. A major unsolved clinical issue is that patients develop chemoresistance, leading to disease progression^10^.

These clinical challenges demand innovative therapies. One conceptually promising strategy is the combination of chemotherapy and targeted drugs such as kinase inhibitors^11–13^. Past advances in the molecular characterization of pancreatic tumors have unveiled key pathways that are critical during disease development and progression^14–16^. Notably, early pancreatic carcinogenesis is driven by oncogenic mutations in KRAS (>90%), leading to aberrant MAPK signaling that dysregulates cell proliferation, survival, and differentiation^17^. Moreover, proteins associated with angiogenesis, insulin signaling, or the AKT/MTOR pathway are frequently overexpressed, possibly also promoting the progression of pancreatic tumors towards more severe phenotypes^18^.

Kinase inhibitors (KI) can effectively inhibit many cancer-related signaling pathways and are important cancer therapeutics today. Currently, at least 30 kinase inhibitors are under clinical investigation for PDAC (phase II or III), either as single agents or in combination with chemotherapy^18^. It is noteworthy that many of these investigated KI are approved drugs in other indications. Such drug repurposing holds the potential to accelerate clinical development by leveraging well-characterized drug safety profiles and pharmacokinetics^19^.

In PDAC, the only approved repurposed targeted therapy in combination with GEM is Erlotinib, a small molecule EGFR inhibitor initially developed for non-small cell lung cancer (NSCLC) ^20^. The approval came as a result of a phase III trial in which the combination displayed a very modest (two weeks) improvement in terms of overall survival over GEM alone in advanced PDAC patients^21^. Further post-approval trials showed that the clinical impact of this drug combination was marginal ^22–24^. In NSCLC, the success of Erlotinib was, in part, due to the response association to certain EGFR mutations, but no such molecular subgroup or predictive response marker could be established for PDAC^25^. Moreover, molecular evidence that this drug combination works in a PDAC-specific context remain sparse^26–28^. We argue that the development of GEM/KI combination therapies in PDAC would immensely benefit from a mechanistic rationale based on specific molecular drug-response biomarkers.

Therefore, in this study, we aimed to discover synergistic combinations of GEM and KI in PDAC and to elucidate the underlying molecular mechanisms of action(s) (MoA) of the observed synergies. First, we screened for drug synergy between GEM and 146 targeted agents (almost all approved and phase III kinase inhibitors to date) in 13 PDAC cell lines. This screen identified the ATR inhibitor Elimusertib to be effective in almost all cell lines, prompting our interest in studying the underlying MoA. Therefore, we characterized the impact of four investigational ATR inhibitors on cellular signaling using a recently introduced dose-resolved phosphoproteomic approach called decryptM^29^. Both phenotypic and molecular datasets are available in ProteomicsDB^30^ for further exploration (https://proteomicsdb.org). Analysis of the proteomic data revealed strong DNA damage induction by GEM and simultaneous suppression of DNA repair pathways as the MoA of the drug combination. These results provide a molecular rationale for trying the combination of GEM and ATR inhibitors in PDAC patients.

## Results

### Phenotypic combination screen of Gemcitabine with molecularly targeted cancer drugs

To systematically identify established drugs that possibly synergize with GEM, we performed a phenotypic drug screen with 146 targeted agents alone as well as in combination with GEM in 12 human PDAC cell lines and one hTERT-immortalized pancreatic duct cell line (HPDE; all 13 are referred to PDAC for simplicity; Fig. 1a, Supplementary Data 1). To facilitate potential clinical translation, we focused on clinically advanced compounds (87 approved, 46 phase III and 13 phase I or II; Supplementary Data 1). Of the 146 drugs, 140 were kinase inhibitors, three PARP1 inhibitors (Olaparib, Talazoparib, Niraparib), and inhibitors of STAT3 (Napabucasin), XPO1 (Selinexor), and SMO (Glasdegib). Cells were treated with 11 doses of the targeted drugs spanning concentrations between 170 pM and 10,000 nM and tested alone or in combination with two fixed concentrations of GEM (GEM low, GEM high; see below), resulting in more than 1,800 pairs of drugs and cell lines. A median z-prime of 0.87 was calculated from both positive and negative controls across the 195 screening plates (384-well format; see methods for details), indicating high data quality and robustness of the screen.

**Fig. 1:**
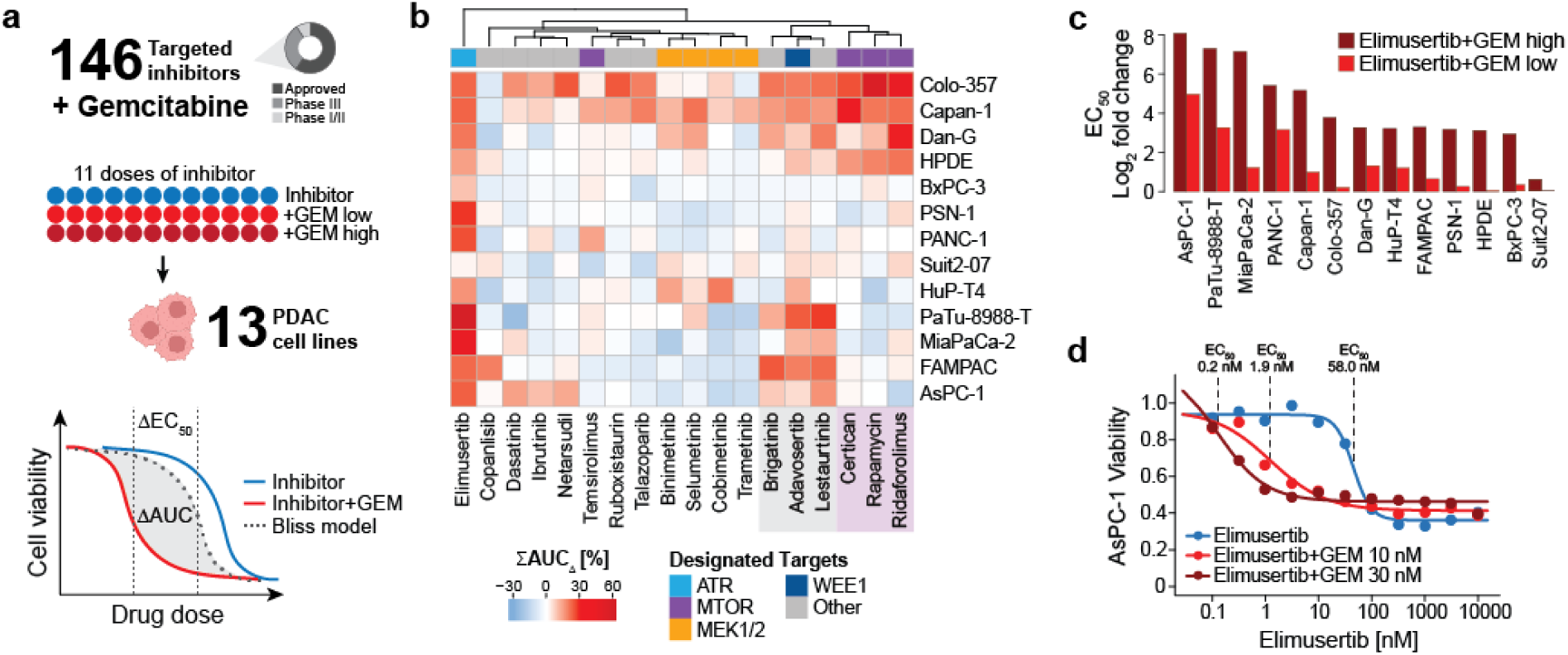
Drug combination screen reveals ATRi Elimusertib to synergize with GEM in most PDAC cell lines. **a** Drug combination screen of 146 clinically relevant inhibitors and GEM in 13 PDAC cell lines. Eleven doses of inhibitor were tested alone or in combination with two doses of chemodrug (GEM high, GEM low), and synergy was assessed using the Bliss model of independence. **b** Heatmap showing the summed shift in AUC (**Σ**AUC_Δ_, in %) for the 18 drugs that synergized with GEM in at least one cell line. A higher **Σ**AUC_Δ_ indicates greater synergy. Drugs are annotated with their designated target proteins. Two clusters of drugs identified from hierarchical clustering are highlighted in grey and purple. **c** Barplot showing the fold change in EC_50_ (log_2_) upon combination of ATR inhibitor elimusertib and GEM high (dark red) or GEM low (light red) for all cell lines. **d** Viability curves upon ATR inhibitor Elimusertib alone (blue) or in combination with either 30 nM GEM (dark red) or 10 nM (light red) in AsPC-1 cells.

In the single-agent screen, 50 of 146 compounds showed efficacy in at least one cell line (area under the curve (AUC) <80%, -log_10_ effective concentration 50 (pEC_50_) >6, and goodness of curve fit of squared Pearson correlation coefficient (R^2^) >0.8; Supplementary Data 2). Eighteen inhibitors were efficacious in more than half of the cell lines (Supplementary Fig. 1a). The two most active drugs were the cell cycle inhibitors Dinaciclib (CDKi) and Volasertib (PLKi), with median AUCs of 40% and 54% across all cell lines, respectively. Most cell lines also responded to the MTOR inhibitor Sapanisertib (median AUC = 54%). Interestingly, inhibition of MTOR by Rapamycin, Everolimus, Temsirolimus and Ridaforolimus showed much less efficacy (median AUC >80%), suggesting off-targets as the cause of the observed efficacy of Sapanisertib. PDAC cells also responded to inhibition of MEK1 and MEK2 (Cobimetinib, Copanlisib, Trametinib) with a median AUC of <70%, and to the SRC inhibitor Tirbanibulin (KX2-391) with a median AUC of 60%. GEM alone generally caused stronger responses than the targeted drugs, with AUCs ranging from 21% to 86% and EC_50_ values of as low as 1 nM (Supplementary Fig. 1b, Supplementary Data 2). Next, we analyzed the drug sensitivity of immortalized duct cell line HPDE to detect effects unrelated to pancreatic cancer. HPDE cells were particularly sensitive towards the CDKi Dinaciclib (AUC = 38%) and also responded to the mTOR inhibitor Sapanisertib, the XPO1 inhibitor Selinexor, and the receptor tyrosine kinase inhibitors (RTKi) Afatinib (EGFR), Neratinib (HER2), and Dacomitinib (EGFR; median AUC <60%).

Based on these results, GEM concentrations for the combination screen were set as follows: 10 nM and 30 nM GEM (GEM low, GEM high) for the four least GEM-sensitive cell lines (PANC-1, AsPC-1, MiaPaCa-2, PaTu-8988-T: EC_50,GEM_ ≳10 nM), and 1 nM and 3 nM GEM (GEM low, GEM high) for all other cell lines (EC_50,GEM_ <10 nM; Supplementary Data 3).

### Most targeted drugs do not synergize with GEM

Combinations of drugs may lead to stronger or weaker phenotypic effects depending on what biology they target. In addition, positive combined effects can be simply additive or synergistic. To estimate if any drug combination yielded a desired synergistic effect, we applied the Bliss model of independence^31^ (see methods for details). In essence, the Bliss model predicts the dose-response curve for a drug combination treatment based on the two single-drug dose-response curves under the assumption that effects are additive. If the experimentally determined dose-response curve of the same combination is more potent than the Bliss model, the effects are termed synergistic (Fig. 1a). For the purpose of this study, the magnitude of synergy was quantified by summing up the observed difference in AUC (**Σ**AUC_Δ_) for the two GEM concentrations. Of all 146 drugs, 18 showed synergy with GEM in at least one cell line (using the following thresholds: **Σ**AUC_Δ_ >10%, and AUC <80%, pEC_50_ >6, R^2^ >0.8 in at least one of the two GEM combinations; Supplementary Fig. 1c, Supplementary Data 2). Clustering of **Σ**AUC_Δ_ values grouped drugs with the same designated targets (Fig. 1b). No synergy was observed in any cell line for the potent cell cycle inhibitors Dinaciclib and Volasertib that showed strong effects as single agents (Supplementary Data 2). Further, we could not detect synergy for GEM and EGFR inhibition even though this drug combination is in clinical use for PDAC patients^20^. Moreover, none of the other 46 tested RTKi had any synergistic effect when combined with GEM (Supplementary Fig. 1c). Although the current panel of cell lines may not be sufficiently representative of all potential PDAC subtypes, this general lack of synergy between RTKi and GEM *in vitro* is in line with the observation that the combination of GEM and EGFRi Erlotinib is also rather ineffective in the clinic^22–24^. This highlights the importance of rational combination treatment design.

### MTOR and WEE1/CHEK1 inhibitors synergize with GEM in certain cellular contexts

Among the 18 synergistic drug combinations were 17 kinase inhibitors. The PARP1-inhibitor Talazoparib showed synergy in two cell lines. Two groups of inhibitors commonly sensitized a few but not all PDAC cell lines, indicating context-dependent drug synergy. For instance, the three MTOR inhibitors Ridaforolimus, Rapamycin, and Everolimus synergized with GEM in Colo-357, Capan-1, and Dan-G cells. We note that these drugs also strongly sensitized normal HPDE cells to GEM from which one may expect increased issues with clinical toxicity. Another three kinase inhibitors chemo-sensitized a larger set of PDAC cell lines, namely the WEE1 inhibitor Adavosertib (7 cell lines), the ALK inhibitor Brigatinib, and the broad-spectrum inhibitor Lestaurtinib (5 cell lines each). Albeit the distinct designated targets of these drugs, their synergistic behavior across multiple cell lines could be caused by a common underlying mechanism, such as off-target effects. To identify candidate off-targets, we mined two large-scale KI target profiling data sets based on the Kinobeads technology^32,33^ (Supplementary Data 4). In both studies, Adavosertib showed high affinity to three kinases (its designated target WEE1, apparent dissociation constant (*K*_D_^app^) = 12 nM; ADK, *K*_D_^app^ = 12 nM; MAP3K4, *K*_D_^app^ = 52 nM). Lestaurtinib and Brigatinib potently interacted with >20 kinases (*K*_D_^app^ <100 nM), including CHEK1 (*K*_D_^app^ = 70 nM). Despite the lack of one single common target protein, we note that CHEK1 and WEE1 both act as key checkpoint regulators in response to DNA damage (as induced by GEM), suggesting that rather the same biological process explains the observed synergy^34^.

### Gemcitabine and ATR inhibitors synergize across almost all PDAC cell lines

While most drugs did not generate synergy in most cell lines, there was one notable exception. The ATR inhibitor (ATRi) Elimusertib showed synergy in 12 of 13 cell lines (Fig. 1b; Supplementary Data 2). The strong chemosensitizing effects manifested in >10-fold improved EC_50_ values in nearly all affected cell lines (Fig. 1c). Interestingly, strongest synergy between Elimusertib and GEM was observed in the four cell lines that were the least sensitive to GEM monotherapy. Potency was most enhanced in AsPC-1 cells, where the EC_50_ for Elimusertib shifted from 58 nM to <2 nM when combined with GEM (Fig. 1d). Of note, immortalized duct cells were also weakly chemosensitized by Elimusertib (**Σ**AUC_Δ_ = 15%), indicating that toxicity may be observed in patients.

We next verified the observed synergy of Elimusertib and GEM in AsPC-1 cells by cell growth assays (Fig. 2a) and including three additional clinical ATRi that were not part of the initial screen: Berzosertib, Ceralasertib, and Gartisertib. As single agents, Elimusertib and Gartisertib were already rather potent (EC_50_ of 113 nM and 128 nM, respectively), while Berzosertib and Ceralasertib were much weaker (EC_50_ of 766 nM and 1,720 nM, respectively; Supplementary Data 5). Then, we turned the drug combination design around and titrated GEM (1 nM to 3,000 nM) in the presence of three sub-EC_50_ doses of the ATR inhibitors and measured the shift in potency compared to GEM alone (EC_50_ of 45 nM, mean of four experiments; Supplementary Data 5). Of the four ATR inhibitors, Elimusertib and Gartisertib sensitized AsPC-1 cells most strongly to GEM (EC_50_ of 6 nM and 7 nM, respectively), followed by Berzosertib and Ceralasertib (EC_50_ of 18 nM and 12 nM, respectively; Fig. 2b; Supplementary Data 5). Importantly, each of the ATR inhibitors sensitized cells to GEM at extremely low doses of GEM (sub-EC_10_). These results demonstrate that the observed synergy of Elimusertib and GEM in the initial PDAC cells screen can also be generalized to other ATR inhibitors and, therefore, underscores the involvement of the DNA repair machinery of the cell via ATR kinase.

**Fig. 2:**
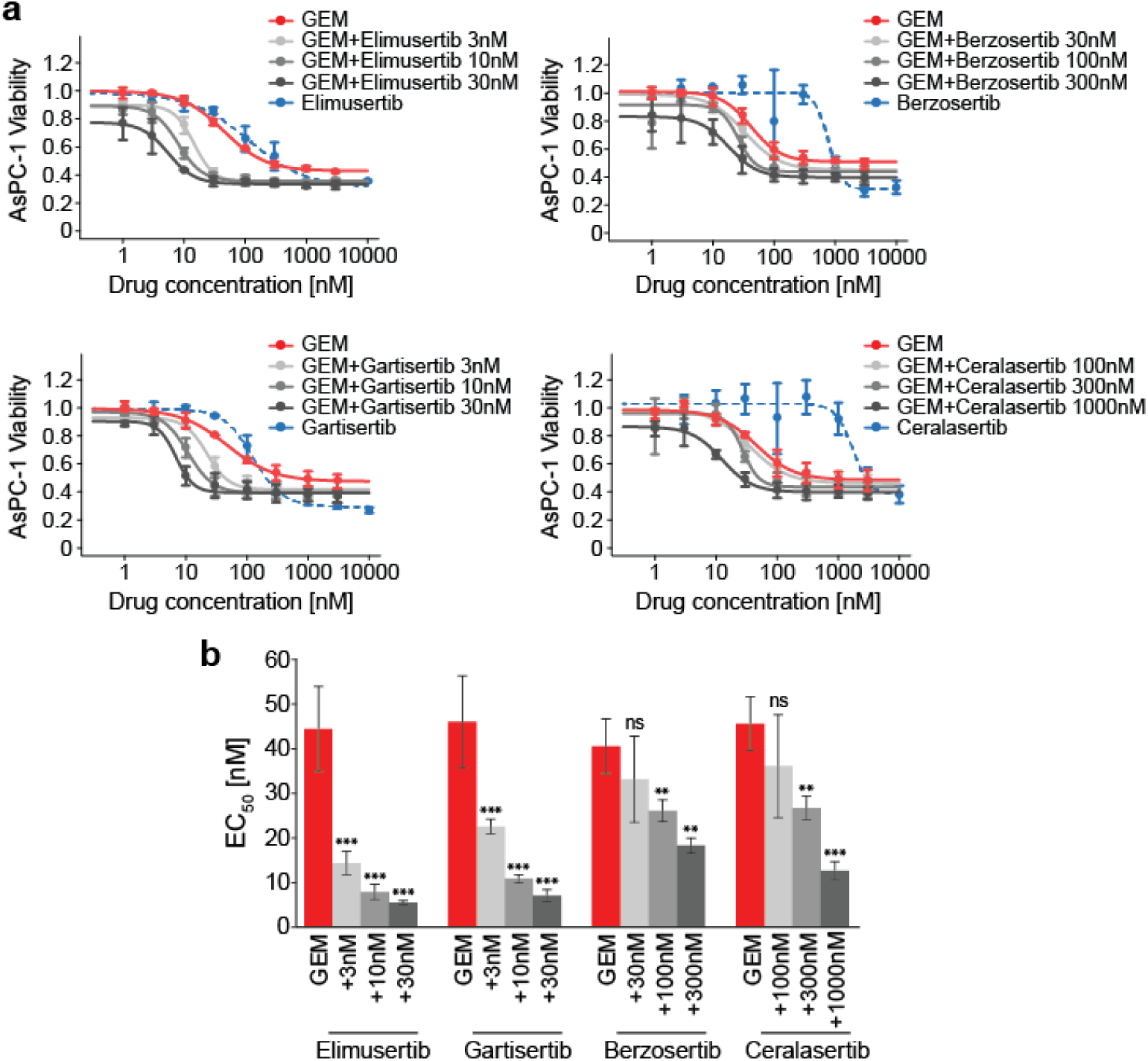
Synergy of Elimusertib and GEM in PDAC cells can be generalized to other ATR inhibitors. **a** AsPC-1 cell viability after treatment with GEM alone (red), GEM in combination with three sub-EC_50_ doses of ATR inhibitor (shades of grey), or ATR inhibitor alone (blue dotted) relative to vehicle. Error bars indicate the +/- standard deviation of triplicates. **b** EC_50_ of GEM alone (red) and GEM in combination with three sub-EC_50_ doses of ATR inhibitor (shades of grey). Error bars indicate the +/- standard deviation of triplicates. Asterisks show the significance level from Student’s t-test against GEM monotherapy (*p*-values were adjusted using the Benjamini-Hochberg procedure; \*\**p*_adj_<0.01; \*\*\**p*_adj_<0.001; ns: non-significant).

### Chemoproteomic target deconvolution reveals the selectivity of clinical ATR inhibitors

The strong chemosensitizing effect of the four tested ATR inhibitors points to the inhibition of ATR kinase activity as the molecular mechanism underlying the observed synergy. Because kinase inhibitors often have more than one target, we subjected Elimusertib and the three investigational ATR drugs Gartisertib, Berzosertib, and Ceralasertib to chemoproteomic target deconvolution in AsPC-1 lysates using the Kinobeads approach^35^ (Fig. 3a) to exclude the effect of an unknown common off-target among the four inhibitors. As expected, all four drugs potently bound ATR kinase and, importantly, ATR was the only common target protein between the four (Fig. 3b; Supplementary Fig. 2a,b; Supplementary Data 6). The Kinobeads data did not reveal any interaction with the structurally and functionally related kinases ATM, DNAPK, or MTOR^36^ (Supplementary Data 6). All four compounds showed off-target binding but to greatly different extents. In contrast to previous findings^37^, we found that Gartisertib is a multi-kinase inhibitor without selectivity for ATR. Indeed, the drug showed stronger interactions with 17 kinases beyond its designated target ATR (*K*_D_^app^ = 318 nM), including GSKA and GSKB (*K*_D_^app^ <10 nM), and several subunits of CSNK2 (*K*_D_^app^ = 10-20 nM). Gartisertib should, therefore, not be applied as a tool for investigating ATR-related cellular effects. Berzosertib also displayed several off-targets in the assay, but ATR was the most potent target by >10-fold, with the exception of ACVR2 (ATR and ACVR2, *K*_D_^app^ <1 nM vs. all others *K*_D_^app^ >10 nM). For Elimusertib (Fig. 3c) and Ceralasertib, only one off-target each was observed, and the affinity to ATR over these off-targets was ∼80-fold (for Elimusertib: ATR, *K*_D_^app^ = 5 nM vs. PIK3C2A, *K*_D_^app^ = 426 nM; for Ceralasertib: ATR, *K*_D_^app^ <1 nM vs. CDK7, *K*_D_^app^ = 84 nM). Together, this verifies that the synergistic phenotypic effect of the four drugs can be explained by the inhibition of ATR.

**Fig 3:**
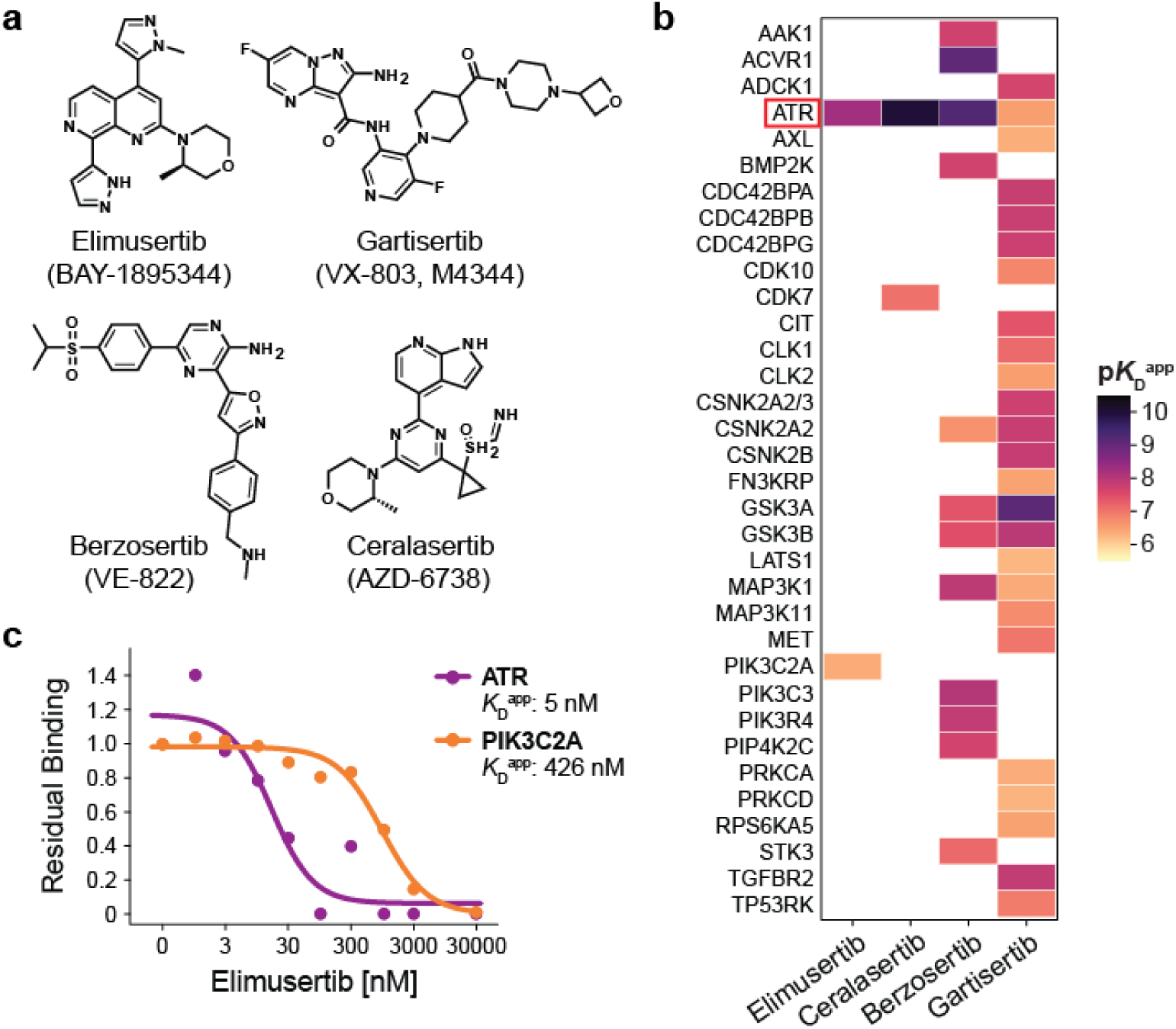
Chemoproteomic selectivity profiling of ATR inhibitors confirms ATR as the only common target kinase. **a** Molecular structures of the four clinical ATR inhibitors Elimusertib, Gartisertib, Berzosertib and Ceralasertib. **b** Heatmap displaying the binding affinities (p*K*_D_^app^) for all target kinases identified with the Kinobeads technology. The designated target kinase ATR is highlighted with a red rectangle. **c** Residual binding of the three kinases ATR, PI3KCB and PIK3C2A on Kinobeads upon increasing doses of Elimusertib. Binding affinities for each target kinase are given as in the legend.

### Clinical ATR inhibitors engage phosphorylation-regulated signaling pathways

To investigate if and how the above ATR inhibitors engage their targets in cells, we followed the decryptM approach^29^ and performed phosphoproteomics on GEM-induced DNA-damaged and subsequently ATRi-treated cells (Supplementary Fig. 3a). Briefly, AsPC-1 cells were pre-incubated with a high dose of GEM (1 µM) for 3 h to induce DNA damage followed by nine doses of ATRi (1 nM – 10 µM) for 1 h (see methods for details). Across all experiments, 25,537 phosphorylated peptides were quantified, of which 20,784 had phosphorylation sites (p-sites) that could be localized with a probability of >0.75 (Supplementary Data 7). Only the latter were used for all subsequent analyses. The resulting ∼64,000 p-site dose-response profiles were statistically evaluated and categorized into significant up- and down-regulated curves by CurveCurator^38^ (see methods for details). As one might expect from the Kinobeads data above, the decryptM analysis uncovered dose-dependent regulation of more p-sites for Berzosertib (n = 840) and Gartisertib (n = 1,015) compared to the more selective compounds Ceralasertib (n = 421) and Elimusertib (n = 525), likely as the result of inhibiting off-targets in cells. Among the significantly inhibited p-sites, there was a striking over-representation of p-sites with a pSQ/pTQ motif (Fig. 4a, Supplementary Fig. 3b), which is known to be the kinase substrate motif of ATR. Other atypical kinases, namely ATM and DNAPK, share the same motif^39^, but they can be excluded as the upstream kinase because they are not targeted by any of the drugs. Among these sites, the well-known ATR substrate and key effector site CHEK1-pS317^40^ was inhibited in a dose-dependent fashion by all four ATRi, which clearly demonstrates reduced kinase activity in cells (Fig. 4b). Inhibition of ATR signaling further manifested in suppressed phosphorylation of the ATR-activator TOPBP1^41^ (pS1504), the homologous recombination mediator BRCA1^42^ (pS1239) as well as FANCD2 (pS319), which is involved in the repair of cross-linked DNA upon phosphorylation by ATR^43^. When summarizing all drug-regulated pSQ/pTQ-sites, Elimusertib and Gartisertib were 10-20-fold more potent in cells (median EC_50_ of 10 nM and 21 nM, respectively) than Ceralasertib and Berzosertib (median EC_50_ of 154 nM and 188 nM, respectively; Fig. 4c). This mirrors the phenotypic cell viability data presented above for the same drugs. We note that the Kinobeads binding affinity data collected in cell lysates would have suggested the opposite trend. This discrepancy may be explained by differences in cellular uptake or export of the different drugs.

**Fig. 4:**
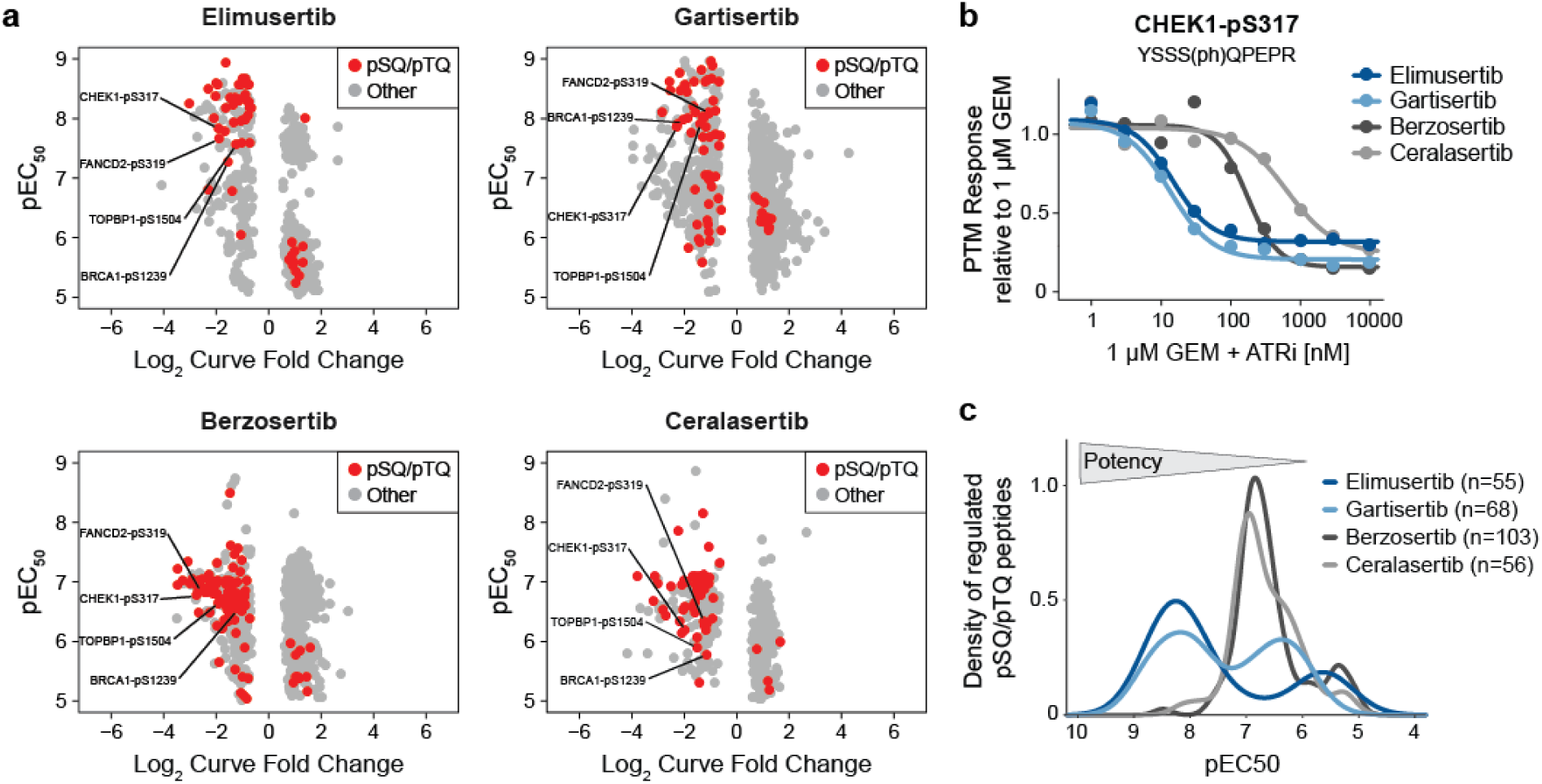
DecryptM reveals cellular engagement of ATR kinase. **a** Potencies (pEC_50_) and curve fold changes (log_2_) from decryptM experiments with Elimusertib and Gartisertib (top) and Berzosertib and Ceralasertib (bottom) in DNA-damaged cells. Each dot represents one dose-response curve of a regulated p-peptide. Red dots indicate phosphorylated peptides containing the pSQ/pTQ motif. Known direct substrates of ATR kinase are annotated by text in the plot. **b** Dose-dependent regulation of CHEK1-pS317 by the four ATR inhibitors in DNA-damaged cells. PTM response was normalized to 1 µM GEM. **c** Density of potencies (pEC_50_) of pSQ/pTQ motif-containing phosphorylated peptides regulated by each of the four ATR inhibitors in DNA-damaged cells. The number of regulated pSQ/pTQ-containing peptides are given in the legend.

Because ATR was confirmed as the only common target of the four inhibitors, we declared p-sites regulated by at least three of the four ATRi as *bona fide* (direct or downstream) ATR-dependent phosphorylation events. This resulted in 298 regulated p-peptides, 42 of which were pSQ/pTQ-sites and thus likely direct substrates of ATR (Supplementary Fig. 3c, Supplementary Data 7). Among the regulated non-pSQ/pTQ-sites were >30 known direct substrates of cyclin-dependent kinases (CDK1 and CDK2; based on kinase-substrate annotations from PhosphoSitePlus^44^; Supplementary Data 7), linking ATR pathway engagement to subsequent impaired cell cycle control. Regulations in this category encompassed p-sites on the cell cycle regulator CHEK1 (pS268), the spindle assembly proteins TPX2 (pT72) and PRC1 (pT481), as well as the proliferation marker MKI67/Ki67 (pT761; Supplementary Data 7).

### ATRi potently block GEM-induced DNA damage response, explaining drug synergy

To pinpoint the cellular mechanism underlying the observed synergy of GEM and ATR inhibition, we took a step back and analyzed the phosphoproteomic data from the angle of which p-sites were regulated by GEM treatment alone (representing induction of DNA damage), and reversed by ATR inhibition in the combination treatment. Treatment of cells with 1 µM GEM alone vs untreated cells revealed 414 statistically significantly GEM-regulated p-peptides (216 up, 198 down) including 135 pSQ/pTQ motif sites (adjusted *p*-value <0.01, log_2_ fold change >1, quantified in at least three out of four replicates; Fig. 5a). As expected, pSQ/pTQ sites were mostly increased upon administration of the DNA-damaging agent and accounted for 62% of all upregulation events (134 out of 216 p-peptides; Supplementary Fig. 4a). ATR inhibition countered 164 of the 416 GEM-induced phosphorylation changes including 36 pSQ/pTQ sites (Fig. 5a, Supplementary Fig. 4b,c). More specifically, ATRi reverted GEM-induced phosphorylation (62 p-peptides including the 36 pSQ/pTQ), and restored GEM-inhibited phosphorylation (102 p-peptides, only non-pSQ/pTQ). Phosphorylation of several CHEK1 p-sites was suppressed upon combination therapy, including the aforementioned CHEK1-pS317 ATR substrate and CHEK1-pS286 (a CDK substrate). The most strongly affected p-site was CHEK1-pS468 (pSQ/pTQ). Phosphorylation of this site was increased >18-fold upon GEM treatment (log_2_ fold change of 4.2, adjusted *p*-value <0.002) and reduced back to baseline levels by three of the four ATRi (not quantified for Gartisertib; Fig. 5b, Supplementary Fig. 4b). Interestingly, despite CHEK1 being a well-known effector protein of ATR, this particular p-site has not yet been studied in depth in the context of ATR signaling^40^.

**Fig. 5:**
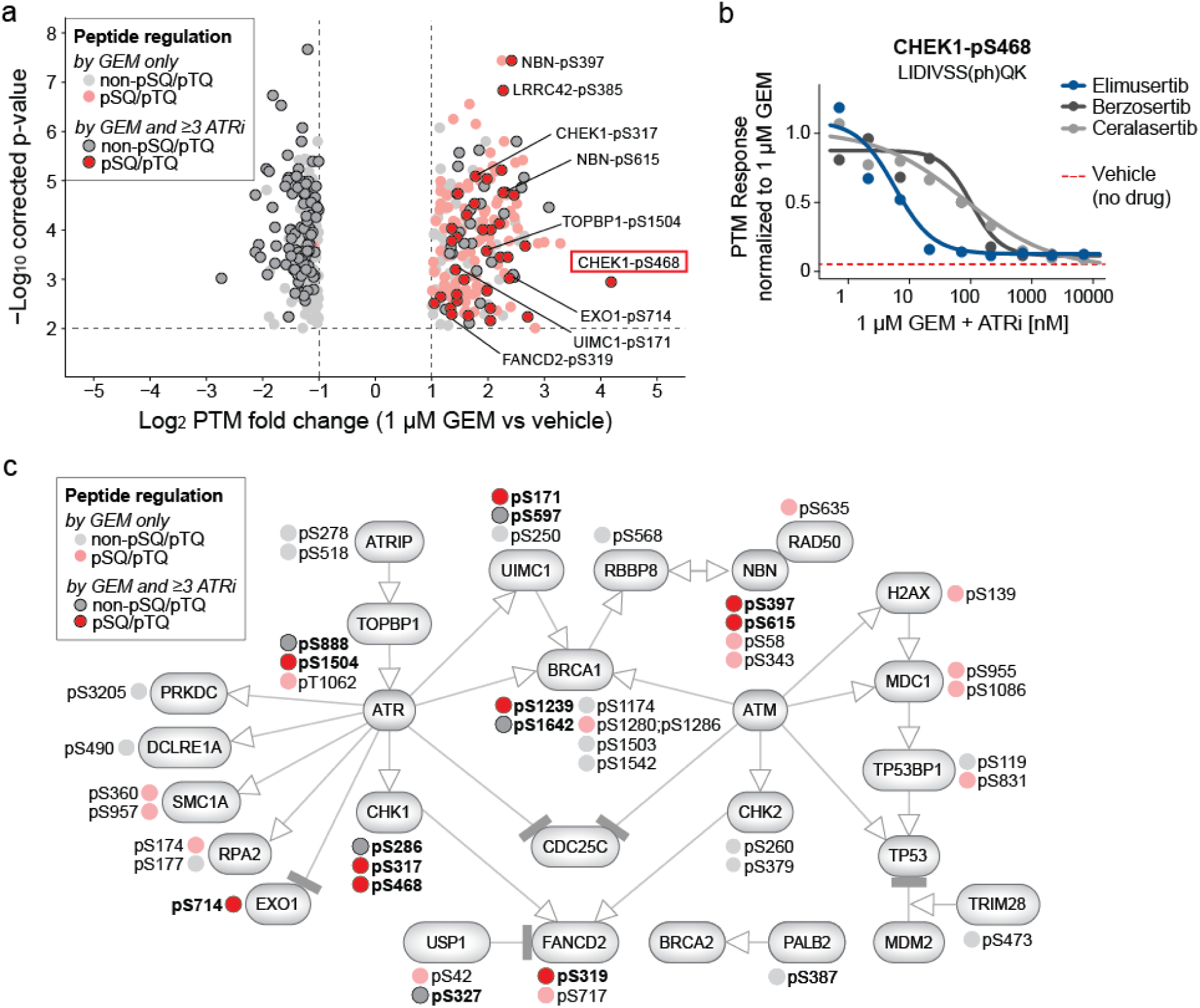
Partial blockage of GEM-induced DNA damage signaling by ATRi explains drug synergy. **a** Volcano plot from t-test of 1 µM GEM vs vehicle after 4 h total treatment (n = 4). *P*-values were corrected for multiple testing using the Benjamini–Hochberg procedure. Phosphorylated peptides were either regulated by GEM but not ATRi (lighter shades), or by GEM and at least three of the four ATRi (darker shades and encircled). Peptides containing the pSQ/pTQ motif are indicated in red. Selected phosphorylated peptides affected upon the combination treatment are annotated by text in the plot, and the site CHEK1-pS468 is highlighted with a red rectangle. **b** Dose-dependent inhibition of CHEK1-pS468 by three of the four ATRi in DNA-damaged cells (not quantified for Gartisertib). PTM response was normalized to 1 µM GEM. Red dotted line indicates the baseline phosphorylation in completely untreated, non-DNA-damaged cells (vehicle normalized to 1 µM GEM). **c** Selected nodes of the WikiPathway WP4016, which was enriched in pathway analysis using PTMNavigator (Score = 14.51). Phosphorylation sites were manually annotated. Phosphorylation sites were either regulated by GEM only (lighter shades), or by GEM and at least three ATRi (darker shades and encircled). Phosphorylated peptides containing the pSQ/pTQ motif are indicated in red, all others are shown in grey.

However, our data suggests that CHEK1-pS468 is a clear marker of drug synergy and may be used as a response biomarker in future investigations in animal studies or clinical trials.

To place our findings in the context of cellular signaling, we performed pathway enrichment and visualization using PTMNavigator^45^. As expected, ATR signaling (WikiPathways entry WP4016) was among the most highly enriched pathways (enrichment score 14.51; Fig. 5c). The proteins annotated in this signaling pathway contained 9 of the 36 ATRi-blocked pSQ/pTQ sites, including the ATR substrates CHEK1 (pS317, pS468), TOPBP1 (pS1504), and FANCD2 (pS319).

Interestingly, a number of proteins and p-sites within this pathway were affected by GEM in an ATR-dependent and ATR-independent fashion. For instance, although GEM-induced DNA damage led to increased phosphorylation of the homologous recombination (HR) protein BRCA1 at seven p-sites, only two of these were reverted by ATRi (pSQ/pTQ pS1239 and non-pSQ/pTQ pS1642; Fig 5c). Other proteins involved in HR-mediated DNA repair were also partially affected by the drug combination (depending on the site). These included the BRCA1 interactor UIMC1 (also known as RAP80) ^46^ and NBN (also known as NBS1), which is essential in the repair of DNA double-strand breaks via HR^47^. Our analysis also revealed proteins which were throughout affected in phosphorylation exclusively by GEM, but not by ATRi on any site. These comprised H2AX (pS139), a well-established DNA damage marker^48^, and additional pSQ/pTQ sites on MDC1 (pS955, pS1086) and TP53BP1 (pS831). These proteins are commonly described as taking part in ATM-mediated DNA repair and cell cycle control^49,50^. Hence, we suspect that these GEM-induced effects arise from ATRi-independent DNA damage signaling, presumably by the other master regulator kinase ATM. Because the four drugs investigated here do not inhibit ATM, these GEM-only p-sites do not relate to the observed drug synergy. This is also in agreement with our observation that ATM inhibition (via AZD-0156) did not synergize with GEM in our drug screen (median **Σ**AUC_Δ_ across all cell lines <2%; Supplementary Table 2). Overall, the above analysis demonstrates the potential of the decryptM approach to discriminate ATR-dependent from ATR– independent signaling in response to GEM-induced DNA damage. It also highlights the importance of p-site level-resolved analysis to understand the mechanism of drug synergy.

The majority of pSQ/pTQ sites we declared to explain drug synergy were not covered by the currently annotated ATR signaling network shown above (>20 sites). These include proteins described to participate in other DNA damage repair processes, and our data suggests a connection to ATR kinase in these pathways as well (Supplementary Data 7). Among them were highly regulated pSQ/pTQ sites on HMGA1 (pS9, pS44), PPM1G (pS201), and TSEN34 (pS136; all >4-fold induced by GEM and inhibited by ATRi; Supplementary Fig. 4b). However, phosphorylation changes also affected proteins that currently lack a clear association to ATR signaling or DNA damage repair. Intriguingly, some of these proteins contain some of the most strongly GEM-induced pSQ/pTQ sites in the dataset: the cytosine methylase NSUN5-pS432 (7-fold), the leucine rich protein LRRC42-pS385 (4.8-fold), and the splicing factor PNISR-pS706 (4.6-fold increase upon GEM and mitigated by at least three ATRi; Supplementary Fig. 4b). Out of these, the strongest combination effect was seen for LRRC42-pS385, which was fully blocked to baseline levels by all four ATRi (Supplementary Fig. 4d). Based on these observations, we conclude that (apart from CHEK1-pS468 mentioned above), these sites may play an important functional role in DNA damage and (potentially non-canonical) ATR signaling, and could serve as drug response markers for the combination of GEM and ATRi.

## Discussion

Because of the very poor prognosis of most PDAC patients, there is a desperate medical need for more successful therapeutic approaches. The combination of standard chemotherapy such as GEM with molecularly targeted drugs may be a valuable strategy but needs a strong molecular rationale to be meaningfully tried in the clinic. Because only few drug combination screens have been performed in the context of PDAC in pre-clinical research settings^51–55^, the current work provides a substantial new molecular resource to provide such a rationale.

One important finding was that drug synergy is rare, which is in line with earlier published studies^51,53,54^. Specifically, the lack of synergy with RTKi *in vitro* indicates that such combinations may only be clinically successful in special and rare cases, raising the need for biomarkers enabling patient stratification.

The second, and more encouraging, key finding was that the ATR inhibitor Elimusertib strongly sensitized almost all PDAC cells to GEM. With hindsight, this may not be unexpected as GEM induces DNA damage, and ATR plays a central role in several aspects of the cellular DNA damage response, notably in detecting and repairing DNA damage during cell replication to ensure genomic stability^56^. In very recently published work, Zhang et al. investigated combined effects of DNA damage response kinase inhibitors, including three ATRi (not Elimusertib), in 62 cancer cell lines covering 12 tumor types^55^. They also found ATRi and GEM to synergize across several cancer cells (including 4 PDAC), and reported ATRi to show stronger overall synergy compared to other DNA damage inhibitors. This is in line with our findings for Elimusertib, and further indicates that the combination of ATRi and GEM may be a promising therapy for other cancer types as well. Although we are not aware of any other large-scale study that systematically assessed drug synergy between ATRi and GEM, others have observed synergy between GEM and inhibitors of CHEK1 and WEE1, two cell cycle regulators that act downstream of ATR in response to DNA damage^34^. For instance, Jaaks et al. tested the combination of 20 targeted drugs and GEM across 30 PDAC cell lines and observed synergy with the CHEK1-inhibitor AZD7762 in nearly half of the cell lines and, to a lesser extent, also with the WEE1 inhibitor Adavosertib^53^. Two further studies reported broad activity of GEM in combination with CHEK inhibition in NSCLC cells^54^, or CHEKi and WEE1i across several cancer entities other than PDAC^51^. In our screen, the WEE1i Adavosertib, as well as Brigatinib and Lestaurtinib, two broad-spectrum inhibitors with off-target activity for CHEK1, also displayed synergy with GEM in nearly half of our PDAC cell line panel. In contrast, the ATR inhibitor Elimusertib synergized with GEM in almost all cell lines. One could, therefore, speculate that the combination of GEM with ATR inhibition may be therapeutically more efficacious in a wider range of heterogeneous PDAC tumors (or other entities) than targeting kinases downstream of ATR. Some evidence for such an interpretation also comes from our single drug screening data, in which the efficacy of Elimusertib was superior to Adavosertib, Brigatinib, or Lestaurtinib.

The dose-dependent phosphoproteomic profiling of Elimusertib following induction of DNA damage by GEM, together with the fact that none of the tested ATR inhibitors showed inhibition of ATM, clearly indicated that the observed drug synergy is rooted in the inability of cells to repair DNA via ATR-dependent mechanisms. For translational purposes, and to be able to link a therapeutic response to the molecular mechanism of a drug, an adequate pharmacodynamic biomarker is required. In our study, we discovered 36 pSQ/pTQ-motif p-sites that could be used to monitor drug response. More specifically, these p-sites were strongly induced by GEM and potently blocked by ATRi. The dataset also confirmed earlier reports that known ATR substrates, including CHEK1, are involved in drug synergy^57,58^. The very robust signal observed for CHEK1-pS468 (pSQ/pTQ; >18-fold induced by GEM and fully countered by ATRi) in the current study makes this p-site particularly promising as a mechanistic marker for the response to the GEM-Elimusertib combination. Interestingly, while CHEK1 is a well-studied protein in DNA damage signaling, very little is known about CHEK1-pS468. In fact, the authors are only aware of a single study which had investigated ATR-dependent phosphorylation of this site some 20 years ago^40^. Another pharmacodynamic biomarker candidate is the methyltransferase NSUN5-pS432 (pSQ/pTQ, 7-fold increase by GEM and neutralized by three ATRi). Although the evidence from the literature is sparse, this site was described to be regulated by ATM kinase in response to DNA double-strand break before^59^. Here, we find that this site is likely a direct substrate of ATR and a promising marker of drug synergy. Finally, another strong marker candidate is LRRC42-pS385 (pSQ/pTQ; ∼5-fold increase by GEM and fully blocked by all four ATRi). LRRC42 is a largely uncharacterized protein. The few cancer-related reports for the protein describe a potential role in lung carcinogenesis and its overexpression in breast tumors, but no connection to DNA damage response signaling has been made yet^60,61^. To the best of our knowledge, phosphorylation of this site has also not been reported yet, let alone its induction by DNA damage or response to drugs. MS-based phosphoproteomics has been used before to study cellular signaling in response to DNA damage induced by radiation or chemotherapy^62–64^. The current work goes substantially beyond the state of the art in several aspects. First, the phenotypic drug screen across many cell lines revealed that ATR inhibition might be a general mechanism by which PDAC cells could be sensitized to GEM. In turn, this may provide a more general way to break GEM resistance in PDAC or other cancers. Second, the study also illustrates that the application of drug dose-dependent chemical proteomic approaches for target identification and selectivity profiling (Kinobeads), as well as pathway engagement measurements (decryptM) greatly facilitates the elucidation of the molecular basis for the observed synergistic drug effects in cells. Third, and aided by the obtained broad (phospho)proteomic coverage, novel ATR substrates could also be identified, including drug response markers that may turn out to be useful for translational studies in the future. In particular, these need to address if the synergy observed for the combination of GEM and ATR inhibitors in 2D cell lines also translates to more sophisticated PDAC models, such as patient-derived xenografts (PDX) or patient-derived organoids (PDO). These experiments will also be valuable to test the suitability of, for instance, the aforementioned CHEK1-pS468 phosphorylation as a patient stratification or therapeutic response marker.

In a recent pre-print, Jadav et al. investigated the MoA of Berzosertib and Gartisertib in US-O2 osteosarcoma cells treated with hydroxyurea^65^. Despite many differences in, for instance, the drug used for induction of DNA damage, the cellular model, and the phosphoproteomic approach applied, individual drug-regulated p-sites are shared between both studies. This partial overlap in the quantified ATR signaling strengthens our confidence in the robustness of our phosphoproteome data. In addition, our work extends the available phosphoproteomic datasets on ATR and DNA damage signaling in cells, and unveils a mechanistic rationale for clinically relevant drug combinations.

Beyond the focus of this report on discovering the basis for the synergy of GEM and Elimusertib, the provided data can be used for several additional purposes. For instance, the dose-dependent single drug screening data was not systematically explored in this report. The same is true for most of the synergistic combinations that affected only a few cell lines. In addition, the many p-sites that were induced by GEM and reverted by ATR inhibition are not yet functionally understood, particularly regarding their role in DNA damage response. For instance, it is possible that some of the ATR inhibitor-regulated pSQ/pTQ p-sites actually arise from ATM activation as a result of ATR inhibition^57^. Eventually, the classification of ATR-independent and ATR-dependent DNA damage response on a single site-resolved level may be explored to better understand the previously reported crosstalk of these kinases^66^.

Notwithstanding these limitations of the current work, our findings provide a mechanistically rational explanation for the combination of GEM and ATR inhibitors in ongoing clinical trials for PDAC^67–69^. In addition, the results also show that Elimusertib produces superior cellular efficacy as a result of the most potent and selective target and pathway engagement among the four investigational ATR inhibitors tested. Hence, one might suggest prioritizing Elimusertib over other drugs for clinical trials investigating GEM-ATRi combinations.

## Methods

### Cell lines and drugs

Cell lines were purchased from ATCC (MiaPaCa-2, AsPC-1, PSN-1, BxPC-3, PANC-1), CLS (FAMPAC, Capan-1), DSMZ (PaTu-8988-T, Dan-G) or creative bioarray (HuP-T4). Cell lines Suit2-07, Colo-357 and HPDE were kindly provided by Prof Kirsten Lauber (LMU, Germany). The identity of all cell lines was authenticated using STR fingerprinting (Suit2-07) or SNP profiling (all others) as provided by Multiplexion GmbH, Germany. Cells were cultivated at 37 °C and 5% CO_2_, and regularly checked to be mycoplasma free. Detailed information on cell culture media is provided in Supplementary Data 1. All media and supplements were purchased from PAN biotech, with the exception of human epidermal growth factor (R&D Systems), bovine pituitary extract (Gibco) and keratinocyte-free medium (Gibco). Only for the drug combination screen, cell media were supplemented with Pen Strep (100 U/mL penicillin and 100 ug/mL streptomycin, Gibco).

Compounds were purchased from different vendors including Selleckchem (Absource Diagnostics GmbH, Germany) and MedChemExpress (Hölzel Diagnostika Handels GmbH, Germany; Supplementary Data 1), and their identity was confirmed by their mass using LC-MS/MS at high resolution. Chemical structures of drugs shown in this work were created using ChemDraw (v23.0.1).

### Drug combination screen

Library drugs were serially diluted from 10 mM to 0.0002 mM in DMSO (11 doses, 1:3 dilution steps) using a Janus Gripper with 384-channel Modular Dispense Technology dispensing head (revvity, previously Perkin Elmer, MA, USA). Afterwards, small amounts were transferred to 384-well intermediate plates (Greiner Bio-One GmbH, Germany). Stock solutions of GEM (1 mM, 3 mM, 10 mM and 30 mM) in DMSO were prepared manually. For drug combination screening, 1,000 to 5,000 cells were seeded into 384-well plates (CulturPlate™, Perkin Elmer, MA, USA) using a Multidrop™ Combi Reagent Dispenser (Thermo Scientific, MA, USA) in 50 µl medium per well 24 h prior to drug treatment (seeding densities see Supplementary Data 1). Compound intermediate plates were filled up with water (1:45.5 dilution) with a Multidrop™ Combi Reagent Dispenser and cells were treated with 2.5 µl of library drug serial dilutions using a Janus Gripper. Then, either 2.5 µl vehicle (for single drug treatments) or 2.5 µl of fixed-dose GEM (for drug combination treatments) were added using a Multidrop™ Combi Reagent Dispenser. The final DMSO concentration in each well was 0.2%. The applied doses of GEM were selected based on the cell lines sensitivity towards GEM as single drug (1 nM and 3 nM GEM for cell lines FAMPAC, Dan-G, Capan-1, Colo-357, PSN-1, HPDE, HuP--T4, BxPC-3, Suit2-07; 10 nM and 30 nM GEM for cell lines PaTu-8988-T, AsPC-1, PANC-1, MiaPaCa-2; see Supplementary Data 3). After 72 h of incubation at 37 °C and 5% CO_2_, viability was measured using the ATPlite 1step Luminescence Assay System according to the manufacturer’s protocol (Perkin Elmer, MA, USA) and the luminescence signals per well were detected with Envision Xcite 2104 plate reader with ultrasensitive luminescence detector (revvity, previously Perkin Elmer, MA, USA). Eight wells per plate without library drug served as negative controls, and sixteen wells per plate containing 5 µM Staurosporine served as positive controls. Control plates not containing any drug were used to correct for incubation effects, and data quality was assessed by calculation of Z-prime of positive and negative controls according to

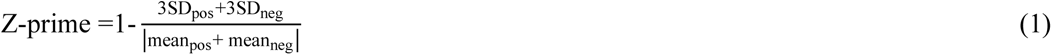

where SD is the standard deviation. All compounds were tested for media solubility with their final dilutions using a NEPHELOstar microplate reader (BMG LABTECH), and a counterscreen was performed to exclude interfering reaction of compounds with ATP instead of cells with ATPlite 1step Luminescence Assay System to detect false active compounds. All data was normalized to vehicle (mean negative controls), and dose-response curves were fitted to a four-parameter log-logistic model, and area-under-curve (AUC, ranging between 0-100%) was calculated as described previously^70^. Single drugs were considered efficacious if AUC <80%, pEC_50_ >6 and R^2^ >0.8.

### Calculation of drug synergy

Synergistic effects were estimated by applying the concept of Bliss Independence^31^. Therefore, a hypothetical dose-response curve was calculated by multiplying the response of the single drugs at each dose used in the combination, describing the expected response when no combination effect occurs, using the following formula:

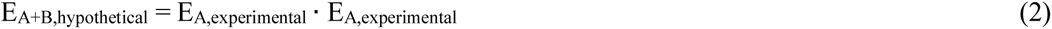

where E_A,experimental_ and E_B,experimental_ are the cell viability upon single-agent treatment with drug A and drug B, respectively, and E_A+B,hypothetical_ is the expected cell viability upon combination. Dose-response curves from the experimental data and the hypothetical data were compared, and shifts in AUC were used to describe combination effects (AUC_Δ_ = AUC_hypothetical_-AUC_experimental_). Drug combinations were defined synergistic if the summed AUC_Δ_ of the two GEM combinations (ΣAUC_Δ_) >10%, and if at least one of the two GEM combinations resulted in AUC <80% and pEC_50_ >6. Only combinations with R^2^ >0.8 in both conditions were considered in the synergy analysis.

### Cell growth assays

Cell proliferation assays of the combination of GEM and clinical ATR inhibitors were performed using the Incucyte live-cell imaging platform (Sartorius). AsPC-1 cells were seeded at 5000 cells/well into 96-well Eppendorf flat bottom plates and allowed to attach overnight. For proliferation assays, cells were treated either with ATRi alone (8 doses, half-logarithmic dilution from 3 nM to 10,000 nM), GEM alone (8 doses, half-logarithmic dilution from 1 nM to 3,000 nM), or a combination of titrated GEM and three fixed, sub-EC_50_ doses of ATRi (0.2% final DMSO concentration; doses see Supplementary Data 5). Cells were incubated for 72 h at 37 °C and 5% CO_2_. To account for pipetting errors, cell confluence readouts at 72 h were normalized to their initial confluence using the Incucyte software. Then, data was normalized to vehicle not containing any drug (n = 5), and dose-response curves were fitted to a four-parametric log-logistic model using the R package *drc*^71^. Shifts in EC_50_ between GEM monotherapy and drug pairs were determined to assess combination effects. Each experiment was performed in triplicates, and data was plotted as mean +/- standard deviation. Significant decrease in EC_50_ (compared to GEM monotherapy) was determined by pairwise Student’s t-test, and *p*-values were adjusted using the Benjamini-Hochberg procedure^72^.

### Chemoproteomic target profiling using Kinobeads

The competition-based drug target profiling assay was performed using Kinobeads ε as described previously^35^. Briefly, 2.5 mg AsPC-1 lysate was incubated with increasing doses of ATRi (1 nM, 3 nM, 10 nM, 30 nM, 100 nM, 300 nM, 1,000 nM, 3,000 nM, and 30,000 nM) or vehicle for 45 min at 4 °C to allow for target binding. This was followed by incubation with Kinobeads ε (17.5 µl of settled beads) for 30 min at 4 °C to enrich kinases. Unbound proteins from the vehicle experiment were collected and subjected to a second pulldown experiment with fresh beads to assess the degree of protein depletion from the lysate by Kinobeads^73^. Enriched proteins were reduced with 50 mM DTT in 8 M Urea, 40 mM Tris-HCl, pH 7.4 for 30 min at room temperature, and alkylated with 55 mM chloroacetamide. After reducing the urea concentration to 1-2 M, proteins were digested with trypsin at 37 °C for 16 h. Subsequently, eluted peptides were desalted using SepPak tC18 µElution plates (Waters), freeze-dried by vacuum centrifugation, and stored at -20 °C.

### LC-MS/MS measurement of Kinobeads experiments

For LC-MS/MS, dried peptides were reconstituted in 0.1% formic acid and analyzed on a Dionex Ultimate3000 nano HPLC (Thermo Scientific) coupled to an Orbitrap Fusion Tribrid mass spectrometer (Thermo Scientific), which was run in data dependent mode. Peptides were first delivered to a trap column (100 µm x 2 cm, packed in-house with Reprosil-Gold C_18_ ODS-3 5 µm resin, Dr. Maisch, Ammerbuch) and subsequently washed with solvent A0 (0.1 % formic acid in HPLC grade water) at 5 µl/min for 10 min. Then, peptides were separated on an analytical column (75 µm x 40 cm, packed in house with Reprosil-Gold C_18_ 3 µm resin, Dr. Maisch, Ammerbuch) at 300 nl/min using a 52 min gradient ranging from 4-32% solvent B (0.1% formic acid, 5% DMSO in acetonitrile) in solvent A1 (0.1% formic acid, 5% DMSO in HPLC grade water). MS1 spectra were acquired in the Orbitrap at a resolution of 60,000 (at m/z 200) over a scan range of 360-1300 m/z using a maximum injection time of 50 ms and an automatic gain control (AGC) target value of 4e^5^ (normalized AGC target 100%). For MS2, up to 12 peptide precursors were isolated (isolation width of 1.2 Th) and subjected to HCD fragmentation using 30% normalized collision energy. Fragmented precursor ions were analyzed in the Orbitrap at a resolution of 30,000, with a maximum injection time of 75 ms and an AGC target value of 1e^5^ (normalized AGC target 200%). The duration of dynamic exclusion was set to 30 s.

### Data processing of Kinobeads experiments

For protein and peptide identification and quantification, raw data was searched against the human reference proteome including isoforms (downloaded from UniProt on March 16, 2021) using MaxQuant^74^ (v1.6.12.0). Oxidized methionine and N-terminal acetylation were set as variable modification, and cysteine carbamidomethylation was set as fixed modification. Label-free quantification (LFQ) and match-between-runs were enabled. All searches were performed with 1% PSM and protein FDR.

Proteins were filtered by potential contaminants, reversed hits and proteins identified only by site. Residual binding of proteins was calculated from LFQ intensities as the ratio between drug doses and vehicle. Data points were fitted using CurveCurator^38^ (default parameters). Kinases were considered a potential target if curve goodness of fit R^2^ >0.7, curve fold change <0.5, curve slope >0.2, and pEC_50_ >6. Only kinases quantified with at least 4 unique peptides in the vehicle experiment were considered. Potential targets were further curated through manual inspection of dose-dependent reduction in MS/MS and unique peptide counts. To calculate apparent dissociation constants (*K*_D_^app^), the EC_50_ value of each protein was multiplied with a correction factor (the protein intensity ratio of the two subsequent pulldown experiments^73^, mean of all four experiments).

### Drug treatment for dose-dependent phosphoproteomics (decryptM)

For decryptM, AsPC-1 cells were seeded at 10 mio cells/10 cm^2^ dish and grown for 48 h before drug treatment (medium was replaced 24 h after seeding). Cells were first pre-incubated with vehicle or 1 µM GEM for 3 h. Then, either vehicle or 9 increasing concentrations of ATRi (1 nM, 3 nM, 10 nM, 30 nM, 100 nM, 300 nM, 1,000 nM, 3,000 nM, and 10,000 nM in medium without fetal bovine serum) were added to cells and incubated for one more hour (final DMSO concentration 0.2%).

### Sample preparation of decryptM experiments

#### SDS cell lysis and protein quantification

Cells were lysed as previously described^75^ with minor differences. Briefly cells were washed twice with PBS and lysed in SDS lysis buffer (2% SDS in 40 mM Tris-HCl, pH 7.6), followed by sonication for 10 min (30s on/30s off) using a Bioruptor (Diagenode). For DNA hydrolysis, samples were boiled at 95 °C for 10 min and incubated with 2% TFA for 1 min, before the reaction was quenched using 4% N-methylmorpholine. Lysate was cleared by centrifugation at 11,000 g for 5 min and protein concentration was determined using the Pierce BCA Protein Assay Kit (Thermo Scientific).

Sample clean-up and digest was performed using the SP3 method on an automated Bravo liquid handling platform (Agilent Technologies, CA, US) as previously described^75,76^ with minor modifications. In brief, 200 µg of protein lysate was mixed with 1,000 µg carboxylate beads (1:1 mix of magnetic SpeedBeads 45152105050250 and 65152105050250, Cytiva) in a 96-well plate. Using 70% ethanol, samples were then precipitated on beads, followed by washing thrice with 80% ethanol and once with 100% acetonitrile. Subsequently, proteins were reduced and alkylated using 10 mM TCEP, 50 mM chloroacetamide, and 2 mM CaCl_2_ in 100 mM EPPS/NaOH, pH 8.5 for 1 h at 37 °C, followed by tryptic digestion at 37 °C for 16 h. Peptides were acidified with 1% TFA and desalted on Chromabond HLB plates (30 µm particle size, 10 mg capacity, Machery-Nagel), using 0.1% TFA for equilibration and sample loading, and 0.1% TFA in 70% acetonitrile for peptide elution. Recovered peptides were freeze-dried by vacuum-centrifugation and stored at -20 °C.

#### TMT labeling

For TMT labeling, dried peptides were reconstituted in 15 µl 100 mM EPPS, pH 8.5, and each treatment condition was labeled with one channel of TMT-11plex reagent (5 µl of 25 µg/µl reagent per reaction, prepared in dry acetonitrile, Thermo Scientific) for 1 h at 23 °C. The reaction was quenched with 0.4% hydroxylamine. Then, samples were pooled, acidified with 1% formic acid, and freeze-dried by vacuum-centrifugation. Labeled peptide pools were desalted on C_18_ Sep-Pak cartridges (37-55 µm particle size, 50 mg capacity, Waters), using 0.1% formic acid as equilibration solvent, and 0.1% formic acid in 60% acetonitrile was used as elution solvent. Eluted peptides were freeze-dried and stored at -20 °C until further use.

#### Offline basic reversed-phase fractionation

Basic reversed-phase (bRP) fractionation of peptides was performed on a Vanquish HPLC (Thermo Scientific). In short, dried peptides were reconstituted in 200 µl 25 mM ammonium bicarbonate, pH 8.0 and directly injected onto a Waters BEH130 XBridge C_18_ column (3.5 µm, 4.6 x 250 mm). At a flow rate of 1000 µl/min, peptides were eluted using a 60 min gradient ranging from 7% to 45% acetonitrile in the constant presence of 2.5 mM ammonium bicarbonate, pH 8.0. Using an automated fraction collector, 96 fractions were collected (between min 7 and 55 after injection). Fractions were acidified with 1% FA and pooled to 48 fractions, before they were freeze-dried and stored at -20 °C.

#### Phosphopeptide enrichment using IMAC

Immobilized metal ion affinity chromatography (IMAC) enrichment of phosphorylated peptides was performed on the Bravo automated liquid handling platform using AssayMAP Fe(III)-NTA cartridges (Agilent). In brief, the 48 dried peptide fractions were reconstituted in 0.1% TFA in 80% acetonitrile and further pooled to 12 fractions (200 µl final volume per well). The phosphopeptide enrichment protocol embedded in the Agilent AssayMAP Bravo Protein Sample Prep Workbench v2.0 software was run. Thereby, the cartridges were first primed with 150 μl 0.1% TFA in acetonitrile at 300 μl/min, and equilibrated with 150 μl 0.1% TFA in 80% acetonitrile at 10 μl/min. Then, pooled fractions were loaded at 5 µl/min and further washed three times with 150 μl 0.1% TFA in 80% acetonitrile at 50 μl/min. Enriched peptides were eluted with 60 μl 1% ammonium hydroxide at 5 μl/min. The collected eluates were acidified with 1% formic acid, freeze-dried and stored at -20 °C until LC-MS/MS measurement.

### LC-MS/MS measurement of decryptM experiments

Phosphorylated TMT-labeled peptides were measured on a Fusion Lumos Tribrid mass spectrometer (Thermo Scientific) coupled to a Dionex UltiMate 3000 RSLCnano System (Thermo Scientific) in data-dependent mode using MS3-quantification. In brief, dried peptides were reconstituted in 50 mM sodium citrate buffer injected onto a trap column (100 µm x 2 cm, packed in-house with Reprosil-Gold C_18_ ODS-3 5 µm resin, Dr. Maisch, Ammerbuch) and subsequently washed with solvent A0 (0.1 % formic acid in HPLC grade water) at a flow rate of 5 µl/min for 10 min. Then, peptides were separated on an analytical column (75 µm x 48 cm, packed in house with Reprosil-Gold C_18_ 3 µm resin, Dr. Maisch, Ammerbuch) at a flow rate of 300 nl/min using an 80 min, two-step gradient. For minutes 0-65, the gradient ranged from 4-22.5% solvent B (0.1% formic acid, 5% DMSO in acetonitrile) in solvent A1 (0.1% formic acid, 5% DMSO in HPLC grade water), and from 22.5-32% solvent B for minutes 65-80. Peptides were ionized using a nano source with 2.1 kV spray voltage. MS1 spectra were acquired in the Orbitrap at a resolution of 60,000 (at 200 m/z) over a scan range of 360 to 1800 m/z, using an AGC target value was set to 4e^5^ and a maximum injection time of 50 ms. The cycle time between consecutive MS1 scans was 3 s. Precursor ions were isolated using a quadrupole isolation window of 0.7 Th, and subjected to CID fragmentation in the linear ion trap using 35% normalized collision energy, with multistage activation enabled. MS spectra were acquired in the Orbitrap at a resolution of 30,000 over an auto scan range, using an AGC target of 5e^4^ and maximum injection time of 60 ms, with the inject-beyond feature enabled. The duration of dynamic exclusion was set to 90 s. Using a charge state-dependent MS3 quadrupole isolation window of 1.2 Th (z = 2), 0.9 Th (z = 3), 0.7 Th (z = 4-6), a new batch of TMT reporter ions was isolated for a consecutive MS3 scan. Using synchronous precursor selection, the top 10 fragment ions of the MS2 scans were isolated and subjected to HCD fragmentation in the linear ion trap using 55% normalized collision energy. MS3 spectra were acquired in the Orbitrap at a resolution of 50,000 over a scan range of 100 to 1000 Th, using an AGC target of 1e^5^ and a maximum injection time of 120 ms.

### Data processing of decryptM experiments

For protein and peptide identification and quantification, raw data was searched against the human reference proteome (downloaded from UniProt on March 16, 2021) using MaxQuant (v1.6.12.0). Oxidized methionine, N-terminal acetylation, and phosphorylation (STY) were set as variable modification, and cysteine carbamidomethylation was set as fixed modification. All other search parameters were kept as default, except for protein FDR which was set to 1. The resulting evidence.txt and msms.txt files were subjected to SIMSI-Transfer^77^ (v0.5.0) using a p10 threshold. Potential contaminants and reversed hits were removed, and only peptides with unambiguously assigned phosphorylation sites with a localization probability of >0.75 were retained. To acquire site-level quantification values, the corrected reporter ion intensities of peptides containing the same phosphosite(s) were summed up. After median centering of corrected TMT reporter channel intensities, channels 1-10 (decryptM in GEM-treated cells) were submitted to CurveCurator (v0.4.1) for curve fitting and classification of significantly regulated dose-response relationships based on relevance score, using a 2D threshold (α-limit: 0.05 and log_2_ fold change limit: 0.45; default parameters^38^). The resulting list of regulated curves was additionally filtered for pEC_50_ >5. Channel 11 served as a vehicle for the chemodrug treatment and was not subjected to curve fitting. Instead, row-wise normalization was performed across all four TMT-batches using the medians of channels 10 and 11. Intensities were then log_2_ transformed and submitted to two-sided t-test analysis in Perseus^78^, with at least three valid values per group (GEM or vehicle) and multiple testing correction using the Benjamini-Hochberg procedure^72^. Regulation was defined significant for corrected *p*-values (*q*-values) <0.01 and absolute log_2_ fold changes >1.

## Data availability

The mass spectrometry proteomics data was deposited to the ProteomeXchange Consortium (https://proteomecentral.proteomexchange.org) via the MassIVE partner repository with the identifier MSV000094340. Processed dose-response data and plots can be downloaded from Zenodo (https://zenodo.org/records/10792252). All phenotypic and proteomic data can be interactively explored in ProteomicsDB (https://proteomicsdb.org).

## Code availability

Data analysis was performed in R version 3.6.3 using publicly-available R packages, or publicly available tools as cited in the respective section. The code for proteomic data analysis and plotting was deposited in a Github repository (https://github.com/kusterlab/Gem_synergy).

## Acknowledgements

We thank Prof Kirsten Lauber from Radiation/Oncology, LMU for providing cell lines Suit2-07, Colo-357 and HPDE, and the Chemical Biology Core Facility at EMBL Heidelberg for providing expertise and support for drug screen experiments. This work was funded by the German Research Foundation (DFG; SFB1321, grant number 329628492), the German Federal Ministry of Education and Research (BMBF; DROP2AI, grant number 031L0305A), and the European Research Council (ERC; TOPAS, grant number 833710). Parts of the figures were created with BioRender.com.

## Author contributions

S.H., J.L. and B.K. designed the research. S.H., L.F. and K.P. performed experiments. S.H. analyzed the data. A.S., M.T., F.P.B., J.M. and F.H. provided assistance in the data analysis. A.S., J.M. and M.T. processed data for ProteomicsDB. S.H. and B.K. wrote the manuscript.

## Supplementary Information

**Supplementary Fig. 1:**
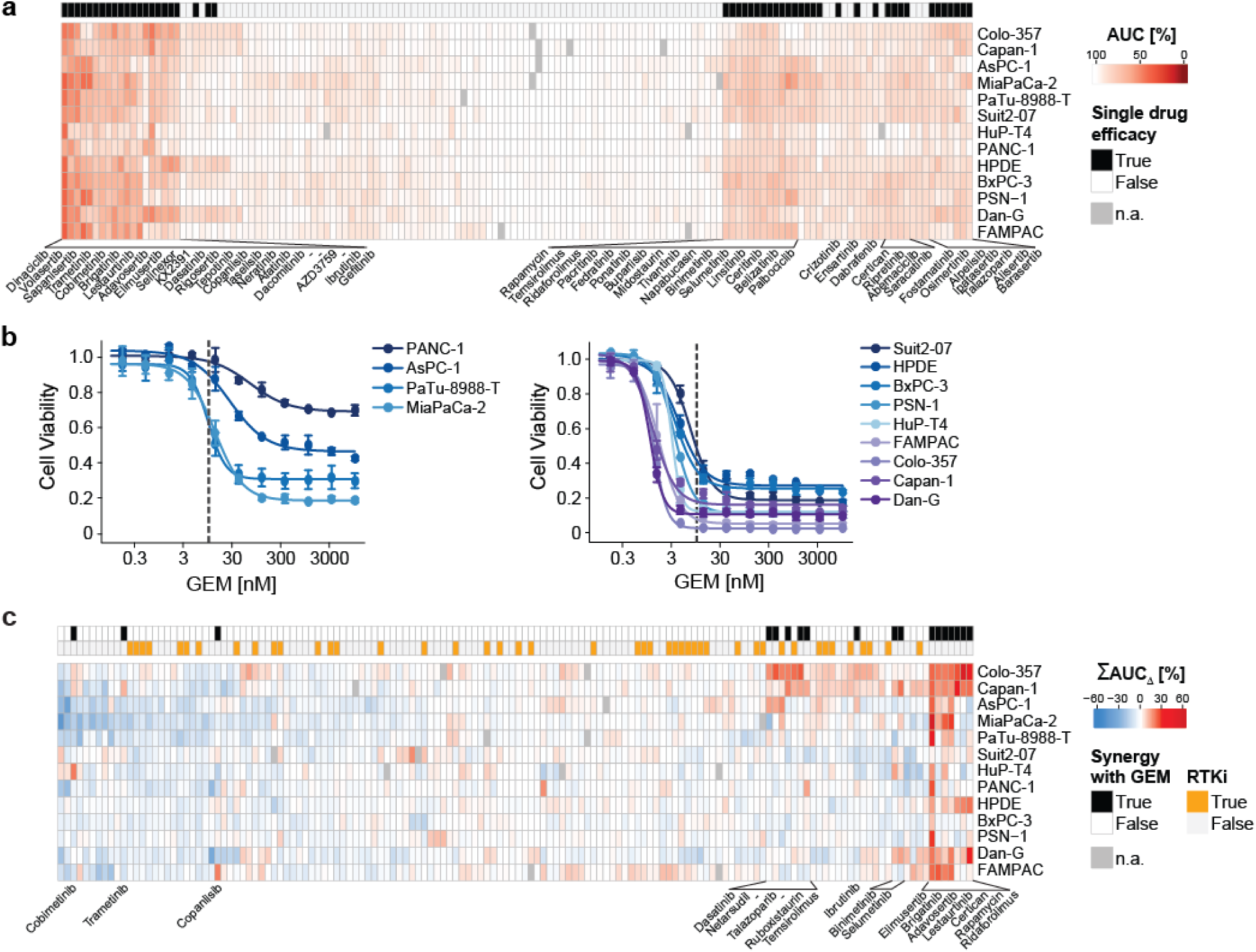
Single drug and combination screening of 146 targeted inhibitors and GEM in 13 PDAC cell lines. **a** Area-under-curve (AUC, in %) of single drug treatments for all 146 inhibitors across 13 PDAC cell lines. A lower AUC indicates greater efficacy. Drugs showing efficacy in at least one cell line are annotated in black, and labeled with text. Missing data (n.a.) is indicated in grey. **b** Cell viability upon treatment with increasing doses of GEM for all cell lines, relative to vehicle. Error bars show the +/- standard deviation of duplicates. Dashed line indicates an EC50 threshold of 10 nM, which was used to separate less sensitive cell lines (left) from more sensitive cell lines (right). **c** Summed shift in AUC (**Σ**AUC_Δ_, in %) upon combination with GEM for all 146 library drugs across all cell lines. A higher **Σ**AUC_Δ_ indicates greater synergy. Drugs showing synergy in at least one cell line are annotated in black, and labeled with text. Inhibitors of receptor tyrosine kinases (RTKi) are annotated in orange. Missing data (n.a.) is indicated in grey.

**Supplementary Fig. 2:**
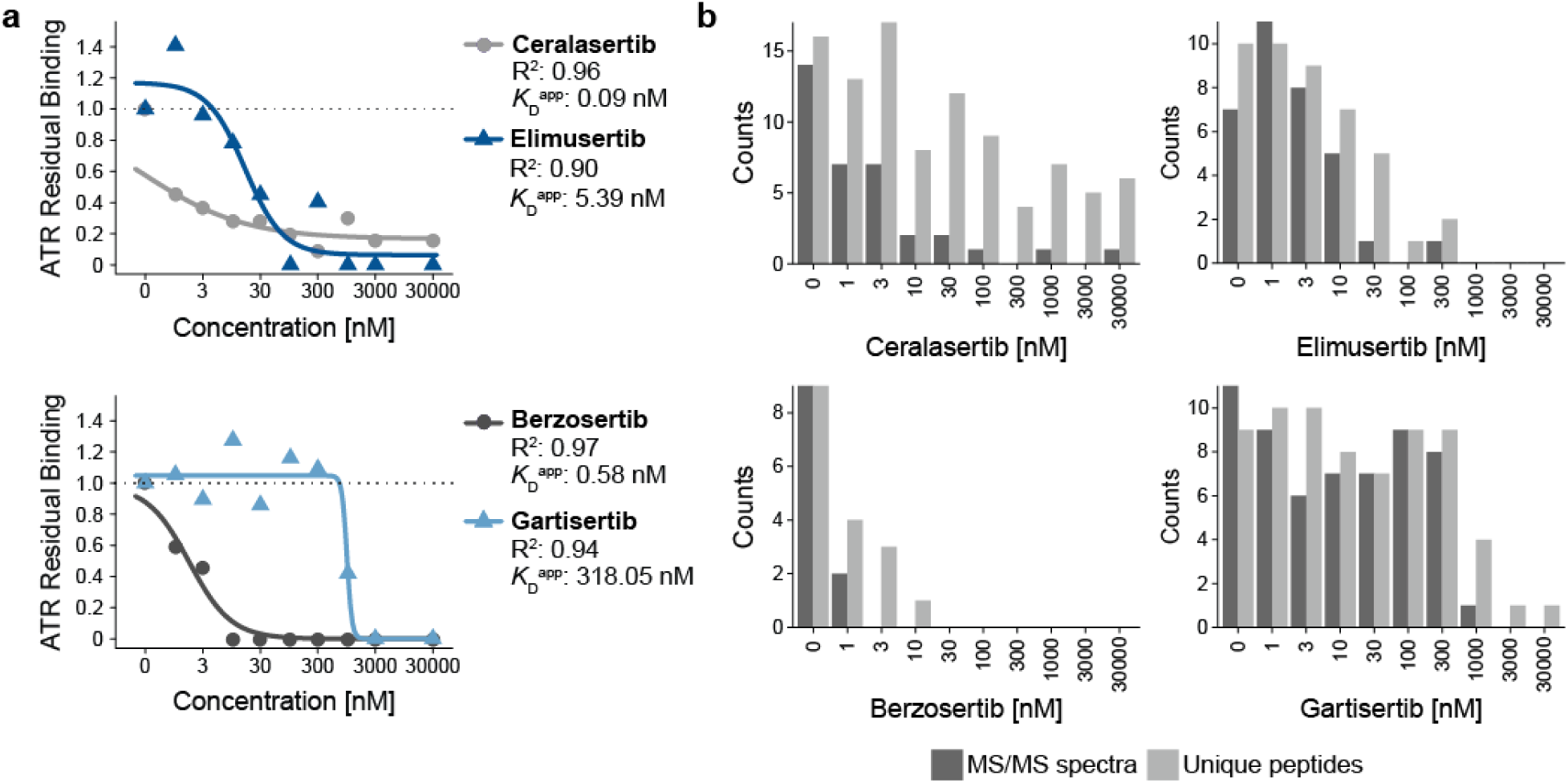
Binding of ATR kinase by clinical inhibitors. **a** Residual binding of ATR on Kinobeads upon increasing doses of Elimusertib and Ceralasertib (top), and Berzosertib and Gartisertib (bottom), based on label-free quantification (LFQ) intensities. Curve fit (R^2^) and apparent affinity constants (*K*D^app^) are given in the legend. **b** Dose-dependent reduction in MS/MS spectra and unique peptides of ATR kinase in pulldown experiment with increasing concentrations of Elimusertib and Ceralasertib (top), and Berzosertib and Gartisertib (bottom).

**Supplementary Fig. 3:**
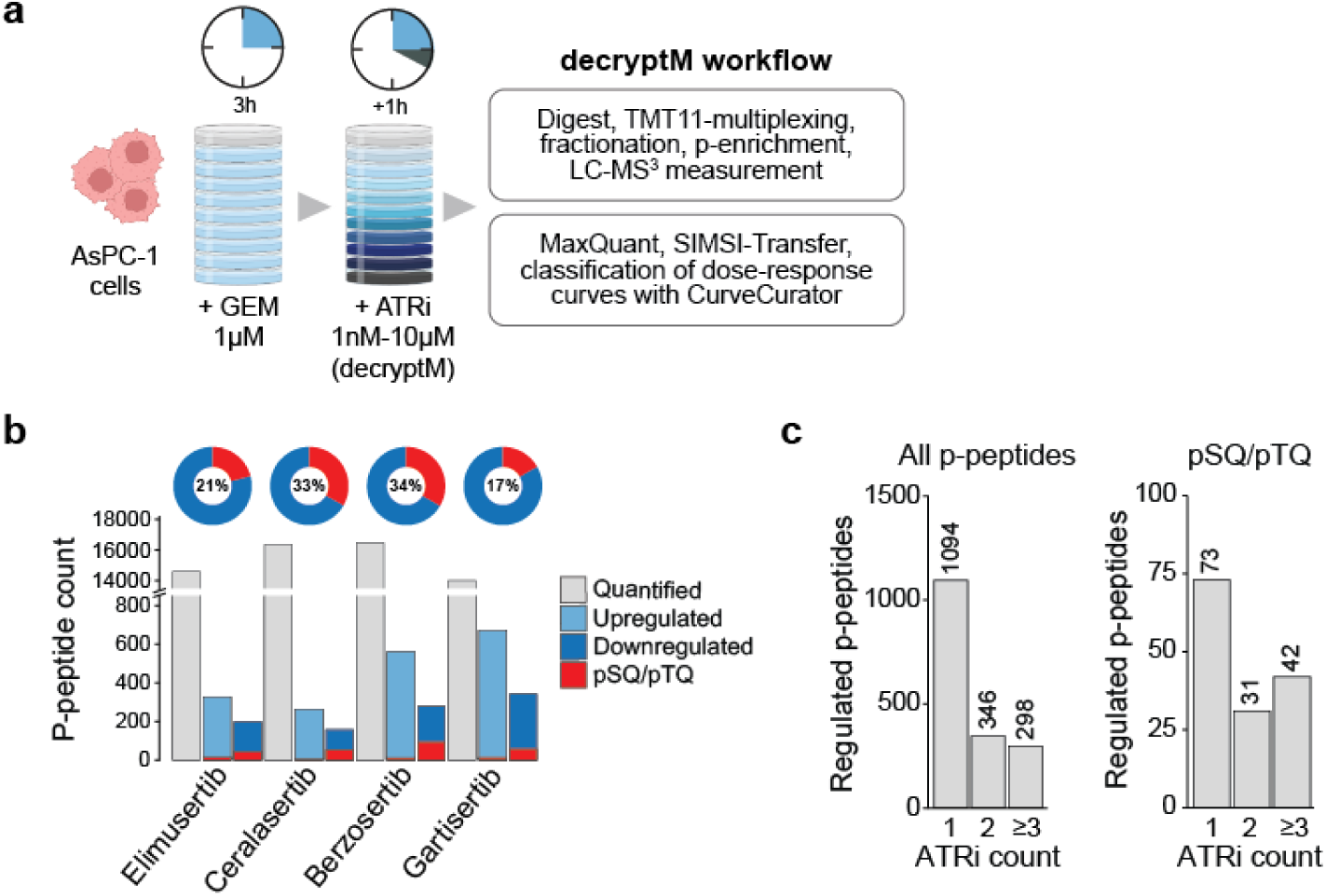
Phosphoproteomic decryptM workflow to study ATR inhibition in DNA-damaged cells. **a** Schematic workflow of decryptM experiments with four clinical ATR inhibitors in DNA-damaged AsPC-1 cells (pre-incubated with GEM). Seeded AsPC-1 cells were incubated with 1 µM GEM for 3 h, followed by nine doses of ATR inhibitor for one additional hour. TMT-labeled, fractionated and phospho-enriched peptides were measured by LC-MS^3^. After data processing with MaxQuant and SIMSI-Transfer, dose-response data was analyzed using CurveCurator. **b** Number of quantified (grey), upregulated (light blue), or downregulated (dark blue) phosphorylated peptides in the four decryptM experiments. Peptides containing the pSQ/pTQ motif are highlighted in red, and the numbers in pie charts indicate the fraction of pSQ/pTQ motif-containing peptides within the down-regulated phosphoproteome. **c** Count of all p-peptides (left) or pSQ/pTQ motif-containing peptides (right) regulated by one, two, or at least three ATR inhibitors in decryptM experiments.

**Supplementary Fig. 4:**
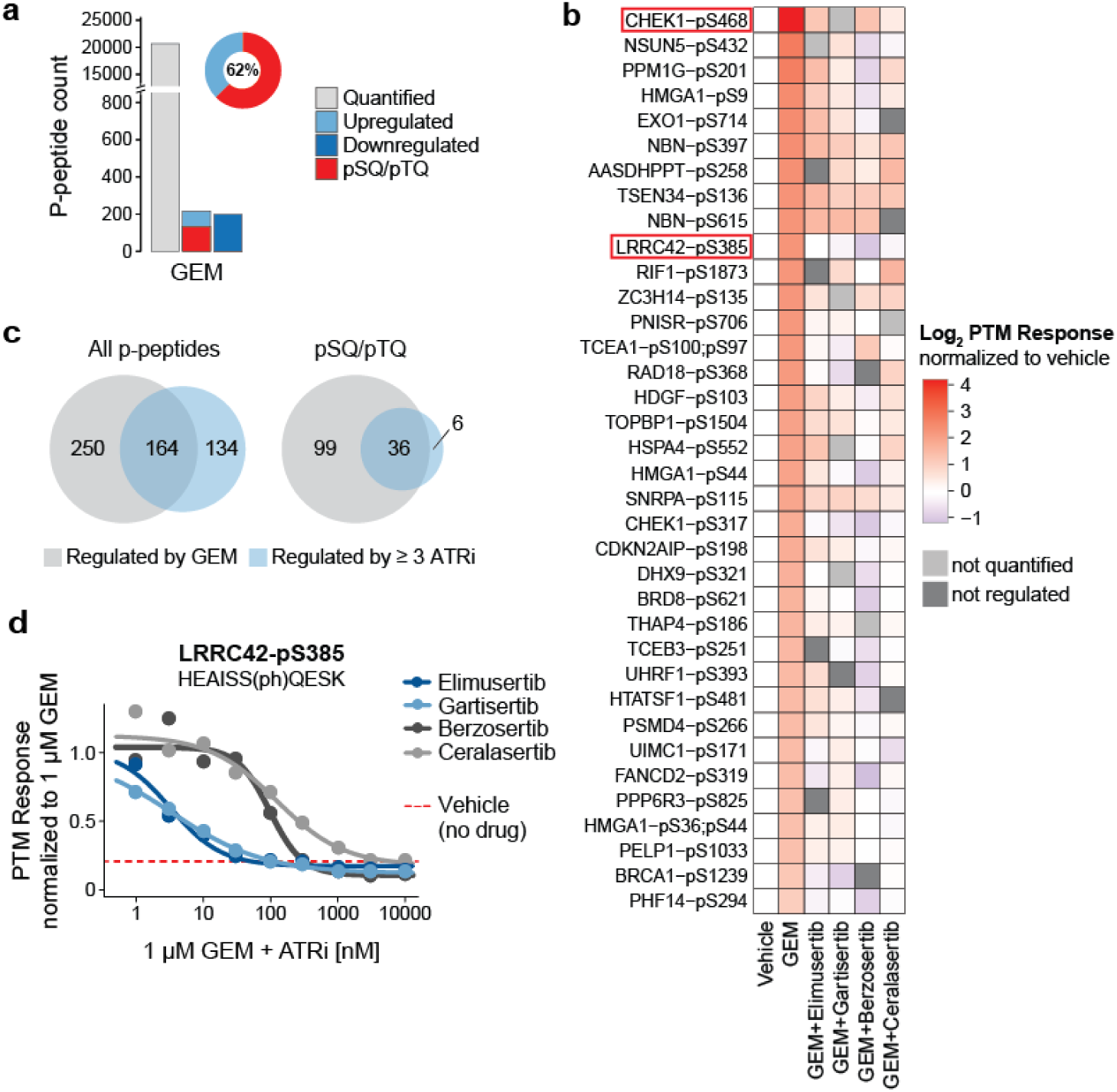
36 GEM-induced pSQ/pTQ sites are counter-regulated by ATRi. **a** Barplot showing the number of all quantified (grey), upregulated (light blue), or downregulated (dark blue) phosphorylated peptides upon treatment of AsPC-1 cells with 1 µM GEM for 4 h (n = 4). Peptides containing the pSQ/pTQ motif are highlighted in red, and numbers in pie charts indicate the fraction of pSQ/pTQ peptides within the up-regulated phosphoproteome. **b** Heatmap of phosphorylation sites induced by GEM and inhibited by at least three of the four ATR inhibitors. Log_2_ fold changes in phosphorylation are shown relative to vehicle (no drug). Light grey indicates missing values (not quantified), and dark grey indicates quantified but not significantly regulated peptides. **c** Overlap in regulated phosphorylated peptides between GEM and at least three out of four ATR inhibitors. Left: all p-peptides, right: pSQ/pTQ motif-containing peptides. **d** Dose-dependent regulation of LRRC42-pS385 by the four ATR inhibitors in DNA-damaged cells. PTM response was normalized to 1 µM GEM. Red line indicates the baseline phosphorylation level in untreated cells (vehicle).

## References

1 Sun, H., Ma, H., Hong, G., Sun, H. & Wang, J. Survival improvement in patients with pancreatic cancer by decade: A period analysis of the SEER database, 1981–2010. Scientific Reports 4, 6747 (2014). 10.1038/srep06747

2 Jemal, A. et al. Annual Report to the Nation on the Status of Cancer, 1975-2009, featuring the burden and trends in human papillomavirus(HPV)-associated cancers and HPV vaccination coverage levels. J Natl Cancer Inst 105, 175–201 (2013). 10.1093/jnci/djs491

3 Siegel, R. L., Miller, K. D., Fuchs, H. E. & Jemal, A. Cancer Statistics, 2021. CA Cancer J Clin 71, 7–33 (2021). 10.3322/caac.21654

4 Park, W., Chawla, A. & O’Reilly, E. M. Pancreatic Cancer: A Review. JAMA 326, 851–862 (2021). 10.1001/jama.2021.13027

5 Burris, H. A., 3rd et al. Improvements in survival and clinical benefit with gemcitabine as first-line therapy for patients with advanced pancreas cancer: a randomized trial. J Clin Oncol 15, 2403–2413 (1997). 10.1200/JCO.1997.15.6.2403

6 Plunkett, W. et al. Gemcitabine: metabolism, mechanisms of action, and self-potentiation. Semin Oncol 22, 3–10 (1995).

7 Kang, J. et al. Nab-paclitaxel plus gemcitabine versus FOLFIRINOX as the first-line chemotherapy for patients with metastatic pancreatic cancer: retrospective analysis. Invest New Drugs 36, 732–741 (2018). 10.1007/s10637-018-0598-5

8 Conroy, T. et al. FOLFIRINOX versus gemcitabine for metastatic pancreatic cancer. N Engl J Med 364, 1817–1825 (2011). 10.1056/NEJMoa1011923

9 Le, N. et al. Real-World Clinical Practice of Intensified Chemotherapies for Metastatic Pancreatic Cancer: Results from a Pan-European Questionnaire Study. Digestion 94, 222–229 (2016). 10.1159/000453257

10 Zeng, S. et al. Chemoresistance in Pancreatic Cancer. Int J Mol Sci 20 (2019). 10.3390/ijms20184504

11 Lei, F., Xi, X., Batra, S. K. & Bronich, T. K. Combination Therapies and Drug Delivery Platforms in Combating Pancreatic Cancer. J Pharmacol Exp Ther 370, 682–694 (2019). 10.1124/jpet.118.255786

12 Garcia-Sampedro, A., Gaggia, G., Ney, A., Mahamed, I. & Acedo, P. The State-of-the-Art of Phase II/III Clinical Trials for Targeted Pancreatic Cancer Therapies. J Clin Med 10 (2021). 10.3390/jcm10040566

13 Lopez, J. S. & Banerji, U. Combine and conquer: challenges for targeted therapy combinations in early phase trials. Nat Rev Clin Oncol 14, 57–66 (2017). 10.1038/nrclinonc.2016.96

14 Waddell, N. et al. Whole genomes redefine the mutational landscape of pancreatic cancer. Nature 518, 495–501 (2015). 10.1038/nature14169

15 Jones, S. et al. Core signaling pathways in human pancreatic cancers revealed by global genomic analyses. Science 321, 1801–1806 (2008). 10.1126/science.1164368

16 Sinkala, M., Mulder, N. & Martin, D. Machine Learning and Network Analyses Reveal Disease Subtypes of Pancreatic Cancer and their Molecular Characteristics. Sci Rep 10, 1212 (2020). 10.1038/s41598-020-58290-2

17 Eser, S., Schnieke, A., Schneider, G. & Saur, D. Oncogenic KRAS signalling in pancreatic cancer. Br J Cancer 111, 817–822 (2014). 10.1038/bjc.2014.215

18 Fang, Y. T., Yang, W. W., Niu, Y. R. & Sun, Y. K. Recent advances in targeted therapy for pancreatic adenocarcinoma. World J Gastrointest Oncol 15, 571–595 (2023). 10.4251/wjgo.v15.i4.571

19 De Lellis, L. et al. Drug Repurposing, an Attractive Strategy in Pancreatic Cancer Treatment: Preclinical and Clinical Updates. Cancers (Basel) 13 (2021). 10.3390/cancers13163946

20 Kelley, R. K. & Ko, A. H. Erlotinib in the treatment of advanced pancreatic cancer. Biologics 2, 83–95 (2008). 10.2147/btt.s1832

21 Moore, M. J. et al. Erlotinib plus gemcitabine compared with gemcitabine alone in patients with advanced pancreatic cancer: a phase III trial of the National Cancer Institute of Canada Clinical Trials Group. J Clin Oncol 25, 1960–1966 (2007). 10.1200/JCO.2006.07.9525

22 Abrams, R. A. et al. Results of the NRG Oncology/RTOG 0848 Adjuvant Chemotherapy Question-Erlotinib+Gemcitabine for Resected Cancer of the Pancreatic Head: A Phase II Randomized Clinical Trial. Am J Clin Oncol 43, 173–179 (2020). 10.1097/COC.0000000000000633

23 Sinn, M. et al. CONKO-005: Adjuvant Chemotherapy With Gemcitabine Plus Erlotinib Versus Gemcitabine Alone in Patients After R0 Resection of Pancreatic Cancer: A Multicenter Randomized Phase III Trial. J Clin Oncol 35, 3330–3337 (2017). 10.1200/JCO.2017.72.6463

24 Biagi, J. J. et al. Adjuvant Chemotherapy and Radiotherapy in Resected Pancreatic Ductal Adenocarcinoma: A Systematic Review and Clinical Practice Guideline. Curr Oncol 30, 6575–6586 (2023). 10.3390/curroncol30070482

25 Tzeng, C. W. et al. Epidermal growth factor receptor (EGFR) is highly conserved in pancreatic cancer. Surgery 141, 464–469 (2007). 10.1016/j.surg.2006.09.009

26 Bartholomeusz, C. et al. Gemcitabine Overcomes Erlotinib Resistance in EGFR-Overexpressing Cancer Cells through Downregulation of Akt. J Cancer 2, 435–442 (2011). 10.7150/jca.2.435

27 Miyabayashi, K. et al. Erlotinib prolongs survival in pancreatic cancer by blocking gemcitabine-induced MAPK signals. Cancer Res 73, 2221–2234 (2013). 10.1158/0008-5472.CAN-12-1453

28 Chen, L. et al. Combination of gemcitabine and erlotinib inhibits recurrent pancreatic cancer growth in mice via the JAK-STAT pathway. Oncol Rep 39, 1081–1089 (2018). 10.3892/or.2018.6198

29 Zecha, J. et al. Decrypting drug actions and protein modifications by dose- and time-resolved proteomics. *Science*, eade3925 (2023). 10.1126/science.ade3925

30 Lautenbacher, L. et al. ProteomicsDB: toward a FAIR open-source resource for life-science research. Nucleic Acids Res 50, D1541–D1552 (2022). 10.1093/nar/gkab1026

31 Bliss, C. I. THE TOXICITY OF POISONS APPLIED JOINTLY1. Annals of Applied Biology 26, 585–615 (1939).

32 Klaeger, S. et al. The target landscape of clinical kinase drugs. Science 358 (2017). 10.1126/science.aan4368

33 Reinecke, M. Identifying small molecule probes for kinases by chemical proteomics, Technische Universität München, (2020).

34 O’Connell, M. J., Raleigh, J. M., Verkade, H. M. & Nurse, P. Chk1 is a wee1 kinase in the G2 DNA damage checkpoint inhibiting cdc2 by Y15 phosphorylation. EMBO J 16, 545–554 (1997). 10.1093/emboj/16.3.545

35 Reinecke, M. et al. Chemoproteomic Selectivity Profiling of PIKK and PI3K Kinase Inhibitors. ACS Chem Biol 14, 655–664 (2019). 10.1021/acschembio.8b01020

36 Blackford, A. N. & Jackson, S. P. ATM, ATR, and DNA-PK: The Trinity at the Heart of the DNA Damage Response. Mol Cell 66, 801–817 (2017). 10.1016/j.molcel.2017.05.015

37 Zenke, F. T. et al. Antitumor activity of M4344, a potent and selective ATR inhibitor, in monotherapy and combination therapy. Cancer Research 79, 369–369 %@ 0008-5472 (2019).

38 Bayer, F. P., Gander, M., Kuster, B. & The, M. CurveCurator: a recalibrated F-statistic to assess, classify, and explore significance of dose-response curves. Nat Commun 14, 7902 (2023). 10.1038/s41467-023-43696-z

39 Kim, S. T., Lim, D. S., Canman, C. E. & Kastan, M. B. Substrate specificities and identification of putative substrates of ATM kinase family members. J Biol Chem 274, 37538–37543 (1999). 10.1074/jbc.274.53.37538

40 Zhao, H. & Piwnica-Worms, H. ATR-mediated checkpoint pathways regulate phosphorylation and activation of human Chk1. Mol Cell Biol 21, 4129–4139 (2001). 10.1128/MCB.21.13.4129-4139.2001

41 Kumagai, A., Lee, J., Yoo, H. Y. & Dunphy, W. G. TopBP1 activates the ATR-ATRIP complex. Cell 124, 943–955 (2006). 10.1016/j.cell.2005.12.041

42 Tibbetts, R. S. et al. Functional interactions between BRCA1 and the checkpoint kinase ATR during genotoxic stress. Genes Dev 14, 2989–3002 (2000). 10.1101/gad.851000

43 Kupculak, M. et al. Phosphorylation by ATR triggers FANCD2 chromatin loading and activates the Fanconi anemia pathway. Cell Rep 42, 112721 (2023). 10.1016/j.celrep.2023.112721

44 Hornbeck, P. V. et al. PhosphoSitePlus, 2014: mutations, PTMs and recalibrations. Nucleic Acids Res 43, D512–520 (2015). 10.1093/nar/gku1267

45 Müller, J., Bayer, F. P., Wilhelm, M., Kuster, B. & The, M. PTMNavigator: Interactive Visualization of Differentially Regulated Post-Translational Modifications in Cellular Signaling Pathways. bioRxiv, 2023.2008.2031.555601 (2023). 10.1101/2023.08.31.555601

46 Yan, J. et al. The ubiquitin-interacting motif containing protein RAP80 interacts with BRCA1 and functions in DNA damage repair response. Cancer Res 67, 6647–6656 (2007). 10.1158/0008-5472.CAN-07-0924

47 Tauchi, H. et al. Nbs1 is essential for DNA repair by homologous recombination in higher vertebrate cells. Nature 420, 93–98 (2002). 10.1038/nature01125

48 Rogakou, E. P., Pilch, D. R., Orr, A. H., Ivanova, V. S. & Bonner, W. M. DNA double-stranded breaks induce histone H2AX phosphorylation on serine 139. J Biol Chem 273, 5858–5868 (1998). 10.1074/jbc.273.10.5858

49 Kim, J. E., Minter-Dykhouse, K. & Chen, J. Signaling networks controlled by the MRN complex and MDC1 during early DNA damage responses. Mol Carcinog 45, 403–408 (2006). 10.1002/mc.20221

50 Stucki, M. & Jackson, S. P. gammaH2AX and MDC1: anchoring the DNA-damage-response machinery to broken chromosomes. DNA Repair (Amst) 5, 534–543 (2006). 10.1016/j.dnarep.2006.01.012

51 O’Neil, J. et al. An Unbiased Oncology Compound Screen to Identify Novel Combination Strategies. Mol Cancer Ther 15, 1155–1162 (2016). 10.1158/1535-7163.MCT-15-0843

52 Falcomata, C. et al. Selective multi-kinase inhibition sensitizes mesenchymal pancreatic cancer to immune checkpoint blockade by remodeling the tumor microenvironment. Nat Cancer 3, 318–336 (2022). 10.1038/s43018-021-00326-1

53 Jaaks, P. et al. Effective drug combinations in breast, colon and pancreatic cancer cells. Nature 603, 166–173 (2022). 10.1038/s41586-022-04437-2

54 Nair, N. U. et al. A landscape of response to drug combinations in non-small cell lung cancer. Nat Commun 14, 3830 (2023). 10.1038/s41467-023-39528-9

55 Zhang, H. et al. Mapping combinatorial drug effects to DNA damage response kinase inhibitors. Nat Commun 14, 8310 (2023). 10.1038/s41467-023-44108-y

56 Cimprich, K. A. & Cortez, D. ATR: an essential regulator of genome integrity. Nature Reviews Molecular Cell Biology 9, 616–627 (2008). 10.1038/nrm2450

57 Wallez, Y. et al. The ATR Inhibitor AZD6738 Synergizes with Gemcitabine In Vitro and In Vivo to Induce Pancreatic Ductal Adenocarcinoma Regression. Mol Cancer Ther 17, 1670–1682 (2018). 10.1158/1535-7163.MCT-18-0010

58 Liu, S. et al. Inhibition of ATR potentiates the cytotoxic effect of gemcitabine on pancreatic cancer cells through enhancement of DNA damage and abrogation of ribonucleotide reductase induction by gemcitabine. Oncol Rep 37, 3377–3386 (2017). 10.3892/or.2017.5580

59 Larsen, D. H. & Stucki, M. Nucleolar responses to DNA double-strand breaks. Nucleic Acids Res 44, 538–544 (2016). 10.1093/nar/gkv1312

60 Fujitomo, T., Daigo, Y., Matsuda, K., Ueda, K. & Nakamura, Y. Identification of a nuclear protein, LRRC42, involved in lung carcinogenesis. Int J Oncol 45, 147–156 (2014). 10.3892/ijo.2014.2418

61 Moody, L., Chen, H. & Pan, Y. X. Considerations for feature selection using gene pairs and applications in large-scale dataset integration, novel oncogene discovery, and interpretable cancer screening. BMC Med Genomics 13, 148 (2020). 10.1186/s12920-020-00778-x

62 Matsuoka, S. et al. ATM and ATR substrate analysis reveals extensive protein networks responsive to DNA damage. Science 316, 1160–1166 (2007). 10.1126/science.1140321

63 Mu, J. J. et al. A proteomic analysis of ataxia telangiectasia-mutated (ATM)/ATM-Rad3-related (ATR) substrates identifies the ubiquitin-proteasome system as a regulator for DNA damage checkpoints. J Biol Chem 282, 17330–17334 (2007). 10.1074/jbc.C700079200

64 Bensimon, A. et al. ATM-dependent and -independent dynamics of the nuclear phosphoproteome after DNA damage. Sci Signal 3, rs3 (2010). 10.1126/scisignal.2001034

65 Jadav, R. et al. Chemo-phosphoproteomic profiling with ATR inhibitors berzosertib and gartisertib uncovers new biomarkers and DNA damage response regulators. bioRxiv, 2023.2004.2003.535285 (2023). 10.1101/2023.04.03.535285

66 Fedak, E. A., Adler, F. R., Abegglen, L. M. & Schiffman, J. D. ATM and ATR activation through crosstalk between DNA damage response pathways. Bulletin of mathematical biology 83, 38 (2021).

67 Middleton, M. R. et al. Phase 1 study of the ATR inhibitor berzosertib (formerly M6620, VX-970) combined with gemcitabine +/- cisplatin in patients with advanced solid tumours. Br J Cancer 125, 510–519 (2021). 10.1038/s41416-021-01405-x

68. 68 US National Library of Medicine. ClinicalTrials.gov, <https://classic.clinicaltrials.gov/ct2/show/NCT04616534> (2020).

69. 69 US National Library of Medicine. ClinicalTrials.gov, <https://clinicaltrials.gov/study/NCT03669601> (2019).

70 Lee, C. Y. et al. Illuminating phenotypic drug responses of sarcoma cells to kinase inhibitors by phosphoproteomics. Mol Syst Biol 20, 28–55 (2024). 10.1038/s44320-023-00004-7

71 Ritz, C., Baty, F., Streibig, J. C. & Gerhard, D. Dose-Response Analysis Using R. PLoS One 10, e0146021 (2015). 10.1371/journal.pone.0146021

72 Benjamini, Y. & Hochberg, Y. Controlling the False Discovery Rate: A Practical and Powerful Approach to Multiple Testing. Journal of the Royal Statistical Society: Series B (Methodological) 57, 289–300 (1995). 10.1111/j.2517-6161.1995.tb02031.x

73 Lemeer, S., Zorgiebel, C., Ruprecht, B., Kohl, K. & Kuster, B. Comparing immobilized kinase inhibitors and covalent ATP probes for proteomic profiling of kinase expression and drug selectivity. J Proteome Res 12, 1723–1731 (2013). 10.1021/pr301073j

74 Cox, J. & Mann, M. MaxQuant enables high peptide identification rates, individualized p.p.b.-range mass accuracies and proteome-wide protein quantification. Nat Biotechnol 26, 1367–1372 (2008). 10.1038/nbt.1511

75 Zecha, J. et al. Data, Reagents, Assays and Merits of Proteomics for SARS-CoV-2 Research and Testing. Mol Cell Proteomics 19, 1503–1522 (2020). 10.1074/mcp.RA120.002164

76 Hughes, C. S. et al. Single-pot, solid-phase-enhanced sample preparation for proteomics experiments. Nat Protoc 14, 68–85 (2019). 10.1038/s41596-018-0082-x

77 Hamood, F., Bayer, F. P., Wilhelm, M., Kuster, B. & The, M. SIMSI-Transfer: Software-Assisted Reduction of Missing Values in Phosphoproteomic and Proteomic Isobaric Labeling Data Using Tandem Mass Spectrum Clustering. Mol Cell Proteomics 21, 100238 (2022). 10.1016/j.mcpro.2022.100238

78 Tyanova, S. et al. The Perseus computational platform for comprehensive analysis of (prote)omics data. Nat Methods 13, 731–740 (2016). 10.1038/nmeth.3901

